# Elucidating Microscopic Events Driven by GTP Hydrolysis Reaction in Ras-GAP System with Semi-reactive Molecular Dynamics Simulation: Alternative Role of Phosphate Binding Loop as Mechanical Energy Storage

**DOI:** 10.1101/2021.05.07.443098

**Authors:** Ikuo Kurisaki, Shigenori Tanaka

## Abstract

ATPase and GTPase have been widely found as chemical energy-mechanical work transducers, whereas the physicochemical mechanisms are not satisfactorily understood. We addressed the problem by examining John Ross’ conjecture that repulsive Coulomb interaction between ADP/GDP and inorganic phosphate (P_i_) does the mechanical work upon the system. We effectively simulated the consequence of GTP hydrolysis reaction in a complex system of Rat sarcoma (Ras) and GTPase activation protein (GAP) in the framework of classical molecular dynamics by switching force field parameters between the reactant and product systems. We then observed *ca.* 5 kcal/mol raise of potential energy about the phosphate-binding loop (P-loop) in Ras protein, indicating that the mechanical work generated via the GTP hydrolysis is converted into the local interaction energy and stored in the P-loop. Interestingly, this local energy storage in the P-loop depends on neither impulsive nor consecutive collisions of GDP and P_i_ with P-loop. Instead, GTP-GDP conversion itself does work on the Ras system, elevating the potential energy. These observations encourage us to challenge a conjecture previously given by Ross. We assert that triphosphate nucleotide hydrolyses do mechanical work by producing emergent steric interaction accompanied with relaxation, namely, a shift of biomolecular system to non-equilibrium state on the reshaped potential energy landscape. Recalling the universality of the P-loop motif among GTPases and ATPases, the observations that we obtained through this study would progress physicochemical understanding for the operating principles of GTP/ATP hydrolysis-driven biological nano-machines.

## Introduction

Various ATPase and GTPase function to regulate cellular processes such as cell mobility^1, 2^, molecular channeling and transporting^3^, protein synthesis^4^, signal transduction^5, 6^ and so on. It has been widely known that the hydrolysis reactions of nucleotide triphosphate are essential steps for their functional expression, while it is not understood satisfactorily from the microscopic viewpoint how such chemical reactions generate mechanical works available for their functional expression.

Knowledge for free energy release of several kcal/mol associated with the hydrolysis has been often remarked as the basis for physicochemical investigation for the molecular mechanism. Nonetheless, it should be noted that this macroscopic, thermodynamic quantity denotes total free energy balance of the reaction, which is obtained from accumulation of any energetic processes (microscopic heat generation, rearrangements of solvation shell and configurational change of biomacromolecules and so on). Thus the value does not necessarily mean how generated work drives molecular dynamics of proteins.

Atomistic characterization of non-equilibrium processes triggered by the hydrolysis reaction should lead to elucidation of microscopic mechanisms for work generation via these hydrolysis reactions. However, this approach appears to be infeasible so far due to experimental difficulty to straightforwardly observe substantially fast, sub-picosecond dynamics with sufficient spatial and temporal resolution.^7, 8^

Molecular simulations are expected as a technical counterpart because of the finer temporal resolution combined with atomic resolution, while it should be noted that chemical reactions are still ‘rare events’ for conventional atomistic simulations. Actually, the reaction times of ATPase and GTPase are usually 1000-fold longer or more than the simulation time length accessible with available computational resources of today^9^.

One example for relevant research is concerned with rat sarcoma (Ras) protein, a monomeric GTPase involved in regulation of cellular signal transduction^10^. The reaction time of GTP hydrolysis by Ras is found around several seconds even under interaction with its own GTPase activation protein (GAP)^11^. It is thus challenging to extend the dynamics of molecular systems to the time domain where such a fairly fast event occurs, by using brute-force MD simulation approaches.

Considering the above circumstances, a feasible approach is to examine a specific physicochemical process, which is important to understand microscopic roles of the reaction process for mechanical work generation and is accessible by conventional molecular dynamic simulations.

It is then worthwhile considering the conjecture by John Ross in 2006, which discusses roles of Coulomb repulsive interaction acting between ADP and inorganic phosphate (P_i_) (or GDP and P_i_) for generation of mechanical work upon the system.^12^ The estimated value of the work in this conjecture is as much as a half of the free energy change via the hydrolysis reaction at maximum so that examining this conjecture appears to give a clue to microscopically understand how the hydrolysis reaction generates mechanical work to drive functional expression of biomacromolecules.

The essence of this conjecture can be found in the two remarks about generation mechanism of mechanical work: (1) impulsive collisions with neighboring chemical groups in proteins by immediate conversion of the repulsive interaction into kinetic energy; (2) consecutive collisions with the neighboring chemical groups with retaining the repulsive interaction for a certain period.^12^ The claims of the conjecture are unambiguous and sound reasonable physicochemically, whereas, to our best knowledge, there are no earlier studies to straightforwardly examine this conjecture by explicitly considering dynamic roles of the repulsive interaction at the atomic level.

In the present study, we examine the two remarks in the above conjecture by theoretically simulating atomistic events driven by hydrolysis reactions of nucleotide triphosphate in protein systems. To verify each of the remarks, we focus on ‘switching force fields in molecular dynamics simulation’ method, referred to as SF2MD hereafter. This kind of methods was already applied to the myoglobin heme-CO dissociation reaction due to technical advantage of SF2MD in simple implementation to existing MD simulation packages and reasonable computational costs comparable to classical MD simulations.^13, 14^ The earlier study succeeded in revealing the microscopic vibrational energy relaxation and the accompanying kinetic energy redistribution.^13^

Then, we extend the earlier SF2MD scheme to be applicable for the complicated reaction process. Hydrolysis of triphosphate nucleotide simultaneously undergoes both bond formation and break, thus being technically more difficult to treat than break of single chemical bond between heme and CO, which was considered in the earlier study.

Then atomistic molecular dynamics simulations with the extended SF2MD are employed to examine GTP hydrolysis reaction of Ras-GTP-GAP system (Figure 1A). We microscopically examine conversion from Coulomb repulsive interaction energy acting between GDP and P_i_ to kinetic energy of these reaction products, evaluate directionality of relative motion between GDP and P_i_ as potential source to do mechanical work and influence of the GTP-GDP conversion on the neighboring functional regions of Ras protein (Figure 1B). Finally, we discuss the essential roles of the hydrolysis reaction by recalling the observed changes of Ras protein.

**Figure 1.**
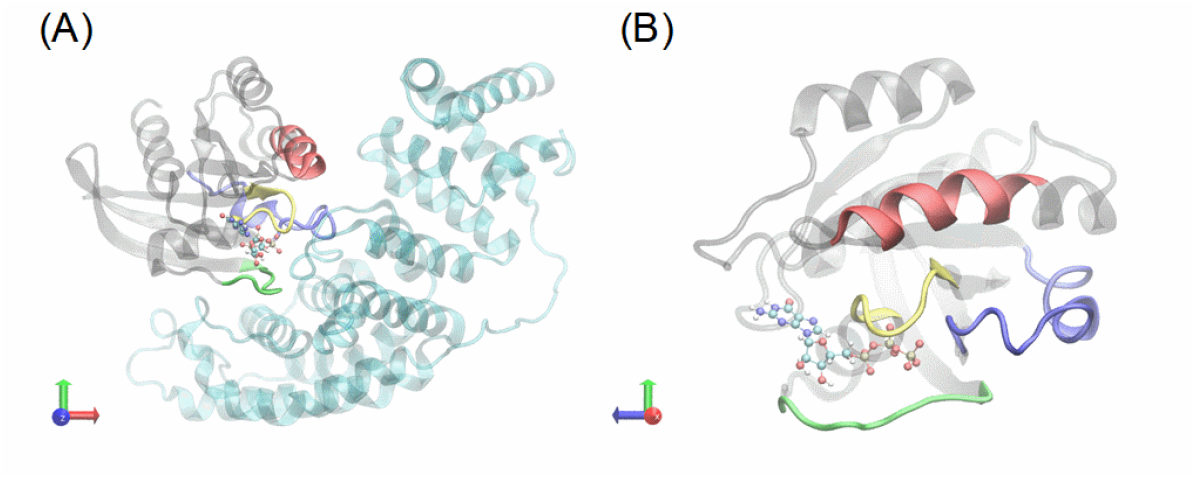
(A) Illustration of Ras-GTP-GAP complex and (B) magnified view of Ras and GTP. Ras and GAP proteins are represented with ribbon, while GTP is drawn by ball and stick. Functional region of Ras is highlighted with different colors as follows, yellow: P-loop; green: Switch-I; blue: Switch-II; red: α-helix 3.

## Materials and Methods

### Simulation setup

To calculate the forces acting among atoms, AMBER force field 14SB^15^, TIP3P water model^16, 17^, and JC ion parameters adjusted for the TIP3P water model^18, 19^ were applied for amino acid residues, water molecules, and ions, respectively. Guanosine triphosphate/diphosphates (GTP/GDP) and Mg^2+^ were described by employing the force field parameters developed by Meagher^20^ and Allnér^21^, respectively.

We here considered doubly protonated inorganic phosphate (referred to as H_2_PO_4_^−^ or Pi, hereafter) as the product of GTP hydrolysis reaction, according to the earlier QM/MM studies^22^. Atomic charges of P_i_ were derived by employing RESP charge calculation with Hartree-Fock/6-31+G* level of theory using Gaussian09^23^ (see **Table S1** and **Figure S1** for details). We selected the level of theory by considering consistency with the GTP and GDP force field parameters^20^. Other force field parameters of P_i_ were derived from general amber force field 2.^24^

All simulations were performed under the periodic boundary condition. Electrostatic interaction was treated by the Particle Mesh Ewald method, where the real space cutoff was set to 9 Å. The vibrational motions associated with hydrogen atoms were frozen by the SHAKE algorithm. The translational center-of-mass motion of the whole system was removed by every 500 steps to keep the whole system around the origin, avoiding the overflow of coordinate information from the MD trajectory format.

### Construction of Ras-GTP-GAP system

We used the X-ray crystallographic structure of Ras-GDP-GAP complex (PDB entry: 1WQ1^25^) to construct a set of atomic coordinates for Ras-GTP-GAP system. Water molecules in the crystal and the bound Mg^2+^ were retained. The atomic coordinates of the aluminum fluoride bound to Ras protein in the vicinity of GDP were employed to give those of γ-phosphate for the GTP system. Nε protonation state was employed for each of histidine residues, and all carboxyl groups in aspartate and glutamate residues were set to the deprotonated state. The one disulfide bond in GAP was formed according to the X-ray crystallography study^25^. The additional details for system construction are described in Supporting Information (*see* **SI 1**).

### MD simulations for structure relaxation

Following temperature relaxation with NVT simulations, 20-ns NPT MD simulations (300 K, 1 bar) were performed and used for structural relaxation. The system temperature and pressure were regulated with Berendsen thermostat^26^ with a 5-ps of coupling constant and Berendsen barostat^26^ with a 0.1-ps coupling constant, respectively. A set of initial atomic velocities were randomly assigned from the Maxwellian distribution at 0.001 K at the beginning of the NVT simulations. The time step of integration was set to 2 fs. This simulation procedure was repeated fifty times by assigning different initial atomic velocities. Each of MD simulations was performed using the Amber 17^27^ GPU-version PMEMD module based on SPFP algorism^28^ with NVIDIA GeForce GTX1080Ti. The further technical details are given in Supporting Information (*see* **SI 2**).

### MD simulation with switching force fields (SF2MD)

Each of 20-ns NPT simulation for structural relaxation is further extended for 106 ps under NVE condition, referred to as front NVE simulation (f-NVE) hereafter. The snapshot structure obtained from the f-NVE is employed to construct Ras-GDP-GAP system by converting GTP and reactant water (W_R_) into GDP and P_i_ (The detailed procedure will be given in the following subsection). The generated Ras-GDP-GAP system is simulated for 106 ps under NVE condition, referred to as back NVE simulation (b-NVE) hereafter. In short, the f-NVE finishes at the switching force field (SF2) point on the overall MD simulation trajectory, and the b-NVE starts from the SF2 point after GTP-GDP conversion.

When we perform the b-NVE, a set of atomic velocities is taken over from the f-NVE except for P_i_ molecule. A velocity vector for each of atoms in P_i_ are set to zero vector. We designed this velocity takeover scheme to perform *seamless* MD simulations with the suppression of thermal noise generation by random reassignment of atomic velocities. Meanwhile, setting the zero velocity to each atom in P_i_ is aimed to circumvent anomalous increase of the kinetic energy via relaxation of P_i_ structure, which may be attributed to inconsistency between atomic velocities given by molecular dynamics of Ras-GTP-GAP system and rearranged atomic configurations as P_i_. A b-NVE is further extended for 20 ns under NVE condition. We also give related remarks on the technical advantage of our SF2MD approaches in mechanical study on triphosphate nucleotide hydrolysis process (**SI 3** and **Figure S2** in Supporting Information).

In b-NVE, atomic coordinates and velocities are recorded by 0.002 ps, 0.05 ps and 1 ps in the first 1 ps, next 5 ps and the last 100 ps, respectively. Finer output of trajectory information for 1-ps NVE simulations is aimed to analyze faster dynamics such as kinetic energy generation found in b-NVE. Symmetrically to b-NVE, in f-NVE, atomic coordinates and velocities are recorded by 1 ps, 0.05 ps and 0.002 ps in the first 100 ps, next 5 ps and the last 1 ps, respectively.

We also performed similar NVE MD simulations for Ras-GTP-GAP system. The f-NVE is simply extended by 106-ps plus 20-ns under NVE condition without GTP-GDP conversion. Although atomic velocities are mostly taken over, those for P_γ_ in GTP and W_R_ were set to zero at the timing of the 106-ps plus 20-ns MD extension. This velocity modification procedure is performed for consistency with simulations for Ras-GDP-GAP system. The 106-ps NVE MD simulation, which is then extended by 20 ns under NVE condition, is referred to as reference NVE simulation (r-NVE), hereafter.

Each 106-ps NVE and the following 20-ns NVE simulations were performed by using the Amber 20^29^ CPU-version PMEMD module and GPU-version PMEMD module based on SPFP algorism^28^ with Tesla V100-PCIE, respectively.

### Rearrangement of atomic coordinates to construct product state

The hydrolysis reaction accompanies both formation and breakage of chemical bonds between reactant molecules. We thus carefully treated such a complicated rearrangement of chemical bonds in contrast to the simple bond breakage discussed in the earlier studies^13, 14^, where we employed QM/MM method to simulate chemical conversion from reactant (GTP + reactant water (W_R_)) to product (GDP + inorganic phosphate (P_i_)). Actually, we found that conventional MM methods did not work to generate physicochemically reasonable P_i_ configuration due to fixed representation of chemical bonds. This technical modification should be useful to avoid anomalous steric crashes among atoms in the product state and following unnatural heat generation unfavorably caused by relaxation of such steric crashes (see **SI 3** in Supporting Information for details). We constructed a Ras-GDP-GAP system by using the snapshot structure obtained from each f-NVE. Positional rearrangement of atoms involved in the GTP hydrolysis reaction (PγO_3_^−^ in GTP and W_R_) was performed by using QM/MM methods with Amber force field 14^15^ and PM3 level of theory^30^. QM/MM simulations were performed by using Amber20 built-in QM/MM interface^31, 32^.

The QM region consists of W_R_ and PγO_3_^−^ in GTP, having 47 atoms in total. In this QM/MM procedure, we deleted the harmonic potential function for the interatomic bond between P_γ_ and Ο2_β_ in GTP by using ParmEd in Amber20^29^. This treatment is technically necessary to define the QM region as closed-shell electronic structure in our simulation with the Amber Package^29^.

By referring to the earlier QM/MM study on GTP hydrolysis in Ras-GAP system^22^, we selected a water molecule as W_R_, which is found in the vicinity of P_γ_ in GTP (a representative configuration will appear later). Besides, we considered the doubly protonated form of inorganic phosphate, H_2_PO_4_^−^, as the reaction product. To obtain atomic coordinates of product state, interatomic distances discussed in the earlier QM/MM study^22^ were constrained by employing harmonic potential with force constant of 500000 kcal/mol/Å^2^ (Table 1). The atom pairs were selected by recalling the earlier QM/MM study. As remarked above, rearrangement of atomic coordinates may cause anomalous increase of kinetic energy if the original velocities are simply taken over. Then, with aiming practically-best seamless connection between f-NVE and b-NVE, we here used the smallest set of interatomic constraints to convert GTP and W_R_ into GDP plus P_i_.

**Table 1.** Pairs of atoms for harmonic potentials in QM/MM simulations and the corresponding equilibrium distances.

| atom 1 | atom 2 | equilibrium distance [$\text{\AA}$] |
| --- | --- | --- |
| $P_\gamma$ in GTP | $O3_\beta$ in GTP | 3.05 |
| $P_\gamma$ in GTP | O in $W_R^\dagger$ | 1.54 |
| $O1_\gamma$ in GTP | $H_1$ in $W_R^\ddagger$ | 0.96 |
| $O1_\beta$ in GTP | $H_1$ in $W_R^\ddagger$ | 1.92 |

$^\dagger W_R$ denotes reactant water molecule whose oxygen atom is found near the $P_\gamma$ in GTP and opposite to the $P_\gamma$ - $O3_\beta$ bond.
$^\ddagger$ One of the two hydrogen atoms in $W_R$ ; the other should be separately referred to as $H_2$ .

Each QM/MM simulation consists of 100 steps of steepest descent method followed by 400 steps of conjugate gradient method. The atomic coordinates except for those consisting of GTP and W_R_ are frozen with ibelly algorithms. Electrostatic interaction was treated by the Particle Mesh Ewald method, where the real space cutoff was set to 9 Å. Interactions between QM and MM regions were considered by electrostatic embedding scheme. The atomic charge in the QM region was set to −1.

### Kinetic energy quenching simulations

The setup for kinetic energy quenching simulations is basically the same as that for the b-NVE. In the kinetic energy absorption scheme, the Ras-GDP-GAP system is combined with the Langevin thermostat with 100-ps^-1^ collision coefficient during the first 1-ps of b-NVE. Meanwhile, the energy minimization scheme employs atomic coordinate optimization for GDP and P_i_ in advance of b-NVE, where MM simulation with 100-step steepest descent method is followed by that with 400-step conjugate gradient method. The other atomic coordinates are frozen with ibelly algorism. A set of atomic velocities except for P_i_ were taken over as in the case of other NVE simulations.

### Analyses of MD trajectories

Energy, hydrogen bond (HB) formation and root mean square deviation (RMSd) were calculated with the cpptraj module in AmberTools 20 package^29^. With regard to each 20-ns NPT MD trajectory, we calculated RMSd to the X-ray crystallography derived Ras-GAP complex structure^25^ using the backbone heavy atoms (i.e., C_α_, N, C and O). RMSd for generated P_i_ structures was calculated for non-hydrogen atoms in P_i_ structure, which was energetically optimized in vacuum with Hartree-Fock/6-31+G* level of theory by using Gaussian 09.^23^ The geometrical criterion of HB formation is as follows: H-X distance was < 3 Å and X-H-Y angle was > 135°, where X, Y and H denote acceptor, donor and hydrogen atoms, respectively.

Mechanical energy was decomposed into three components, namely bonded energy (bond, angle, and dihedral terms), nonbonded one (electrostatic, van der Waals, 1-4 electrostatic and 1-4 van der Waals terms) and kinetic one. Nonbonded energy was calculated under free-boundary condition without distance cutoff by using *energy* command in the cpptraj module implemented in AmberTools 20 package^29^.

The relative orientation between atomic velocities is evaluated by employing angular correlation coefficient, *C_ij_*:

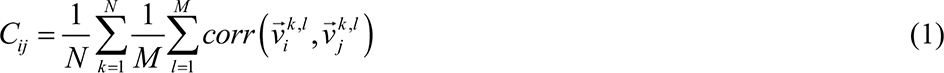

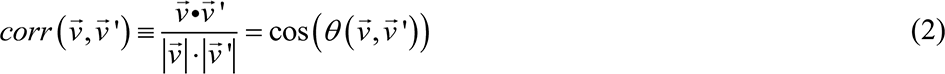

In **Eq. 1**, the indexes *i* and *j* are annotations for specific atoms, while *k* and *l* are identifiers of each independent MD simulation and snapshot structure in the MD trajectory, respectively. Then 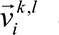 denotes atomic velocity of the atom annotated by *i* at the *k*^th^ snapshot structure in the *l*^th^ MD simulation. The large and small dots in **Eq. 2** mean inner product and multiplication, respectively. In particular, a set of *C_ij_*, 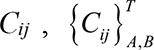, is discussed, where *i* and *j* belong to specific subsets of the system, *A* and *B*, respectively. Snapshot structures are obtained from the time domain, *T*. 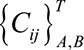 is then represented as frequency distributions or descriptive statistic values, namely, an average and confidence interval.

Molecular structures were illustrated using Visual Molecular Dynamics (VMD).^33^ Error bars are calculated from standard error and indicate 95% confidence interval if there is no annotation.

## Results and discussion

### QM/MM simulations convert GTP with reactant water molecule into GDP and inorganic phosphate

We performed 50 independent unbiased 20-ns NPT-MD simulations of Ras-GTP-GAP to obtain statistically dependable results. Considering temporal change of RMSd (Figure 2), we suppose that Ras-GAP complex reaches structural equilibrium by 20 ns. We then extended each MD trajectory by 106 ps under NVE condition. Each set of atomic coordinates with atomic velocities obtained from the NVE simulation (referred to as f-NVE) was considered to construct a set of atomic coordinates with atomic velocities of the Ras-GDP-GAP system.

**Figure 2.**
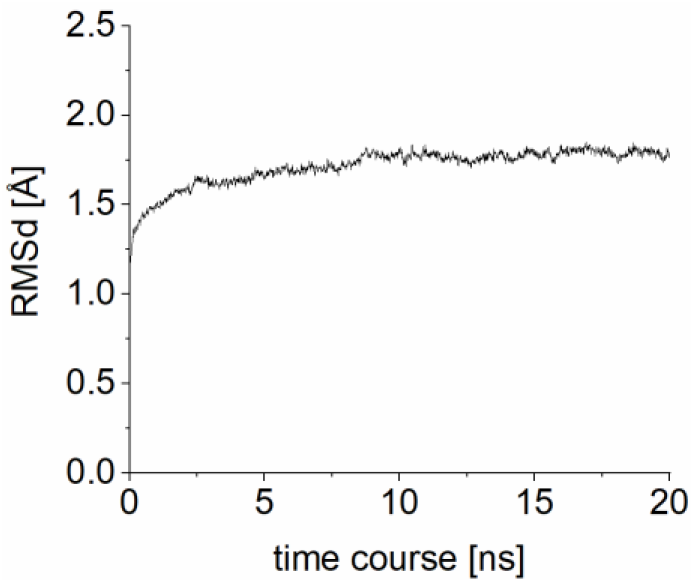
Time course change of root mean square deviation (RMSd) for the Ras-GTP-GAP system. RMSd value at each time point is the average over the 50 independent simulations.

Prior to GTP-GDP conversion by QM/MM simulations, we examined for each of the 50 snapshot structures whether the catalytic metal ion (Mg^2+^) and a reactant water molecule (W_R_) are coordinated near the GTP by recalling the results by the earlier QM/MM study^22^. In each of the 50 snapshot structures, Mg^2+^ is found around P_γ_O_3_^−^ and P_β_O_3_ ^−^ in GTP (a representative structure is given in Figure 3A). Meanwhile, one reactant water molecule was found in the vicinity of P_γ_ in GTP for the 49 snapshot structures. The remaining one snapshot structure missed such a water involved in the hydrolysis reaction, thus being excluded in the following discussion. We finally obtained 49 snapshot structures to construct atomic coordinates of Ras-GDP-GAP system.

**Figure 3.**
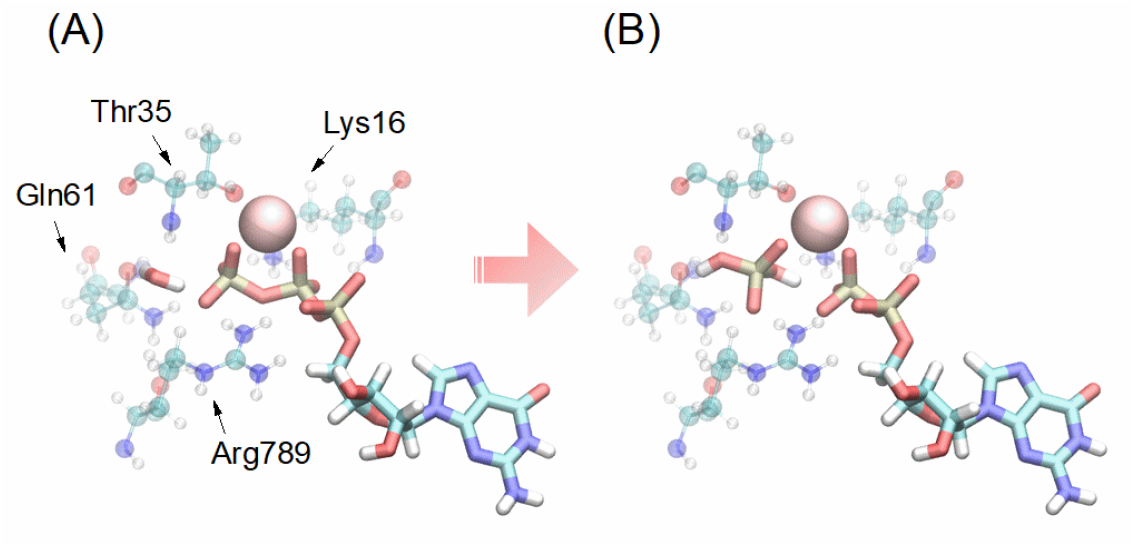
Example of conversion from (A) Ras-GTP-GAP system into (B) Ras-GDP-GAP system by employing the QM/MM simulation. GTP, GDP, H_2_PO_4_^−^ and reactant water molecule are illustrated by solid sticks. Pink sphere is for Mg^2+^ ion. Catalytic residues (Lys16, Thr35 and Gln61 in Ras; Arg789 in GAP) are shown by transparent ball and stick.

We carried out a QM/MM simulation for each of the 49 snapshot structures and converted GTP and W_R_ into GDP and P_i_ (a representative structure is given in Figure 3B). RMSd for P_i_ to the P_i_ structure, which was energetically optimized in vacuum with Hartree-Fock/6-31+G* level of theory, is 0.21 ± 0.02 [Å], denoting that the molecular configuration is finely generated by our QM/MM simulation (**SI 3** in supporting information further discusses technical advantage of the QM/MM-based modeling). Atomic velocities except for those of P_i_ were taken over from the 106-ps NVE-MD simulations for Ras-GTP-GAP system (see Materials and Methods for reason for setting atomic velocities of P_i_ to zero). Each of the 49 sets of atomic coordinates with atomic velocities for Ras-GDP-GAP system was used for the following unbiased 106-ps NVE MD simulations (b-NVE).

### Kinetic energy is increased via relaxation of repulsive Coulomb interaction acting between GDP and P_i_ but promptly loses the coherent directionality of velocities

We analyzed three energy components, kinetic, bonded and nonbonded energy terms, with regard to GDP with P_i_ and tested whether our SF2MD simulations can illustrate the mechanical processes in the conjecture^12^, generation of repulsive Coulomb interaction between them, and the conversion into kinetic energy.

Instantly, we can find apparent increase of the kinetic energy of GDP with P_i_ within the first 0.1 ps (Figure 4A). The increase is partly brought about by relaxation of the repulsive Coulomb interaction between GDP and P_i_, while we will explain the details later. The kinetic energy starts to relax at around 0.1 ps and finally reaches the equilibrium within several picoseconds. Recalling the timescale of energy relaxation, we can say that the change caused by GTP-GDP conversion is relatively fast processes in the context of biomacromolecular dynamics.

**Figure 4.**
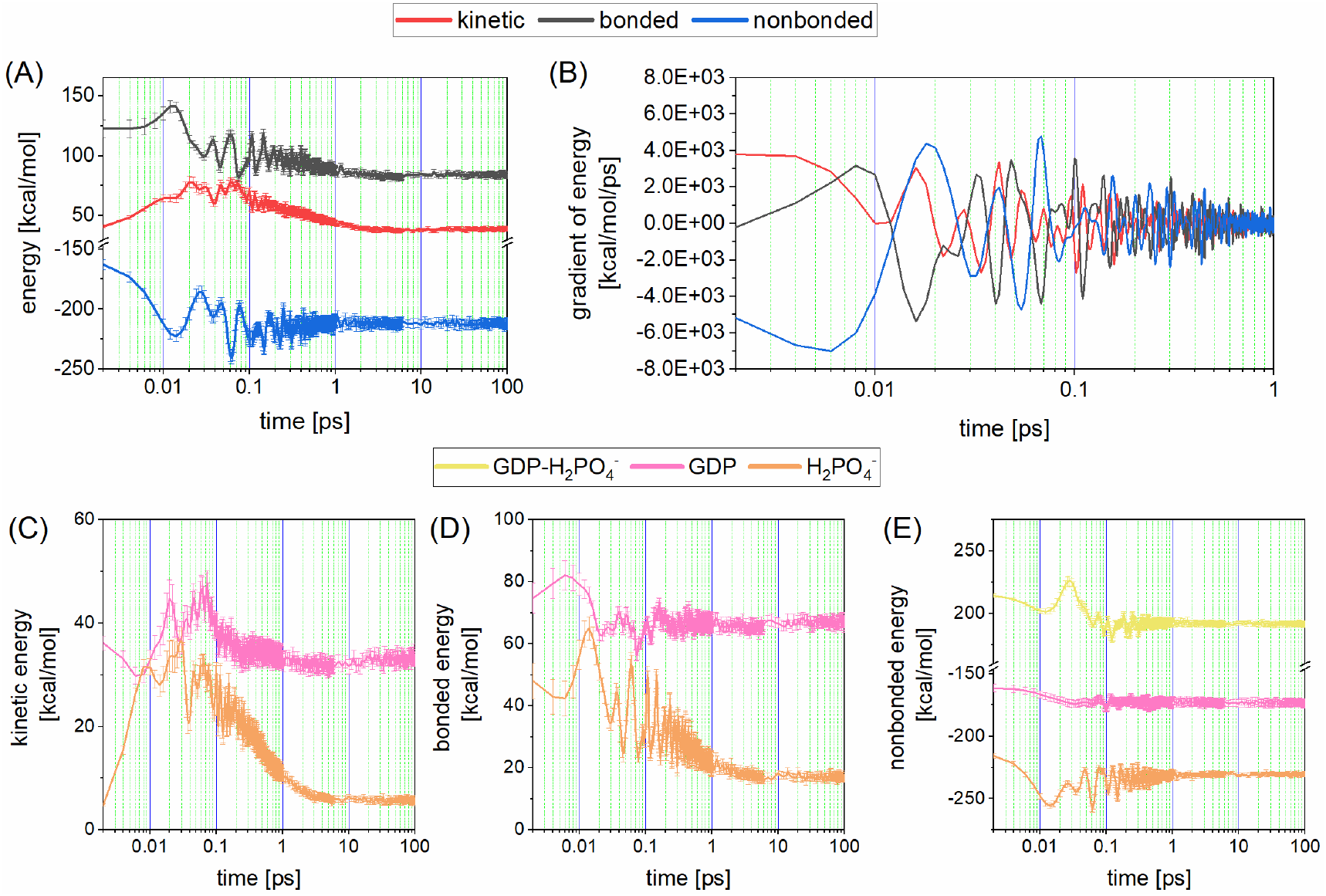
Time course analyses of mechanical energy for Ras-GDP-GAP system. (A) mechanical energy of GDP plus H_2_PO_4_^−^ (inorganic phosphate: P_i_) and (B) the time derivative at each time point, where kinetic, bonded and nonbonded energies were shown with red, black and blue lines, respectively. (C)-(E) Energies of GDP and P_i_; kinetic, bonded and nonbonded energies were for (C), (D) and (E), respectively. Intra GDP, intra P_i_ and inter GDP-P_i_ energies are shown with pink, orange and yellow lines, respectively. An error bar indicates 95% confidential interval.

The increased kinetic energy above is brought about by GTP-GDP conversion. We examined the reference system, where each of 106-ps NVE simulations for the Ras-GTP-GAP system, f-NVE, simply continued under NVE condition without switching force fields. As shown in **Figure S3**, apparent raise of kinetic energy was not observed for this reference system. This result indicates that GTP-GDP conversion is essential to bring about the energy relaxation processes as illustrated in Figure 4. Then, as follows, we examined the microscopic mechanisms of the raise of kinetic energy in the Ras-GDP-GAP system.

The remarkable increase of kinetic energy occurs within the first 0.01 ps, which is accompanied by relaxation of nonbonded energy of GDP with P_i_ (Figure 4A and B). We further examined mechanical energies for each of GDP and P_i_, individually. The increase of kinetic energy, which occurs by the first 0.01 ps, is attributed to relaxation of nonbonded energy for P_i_ (Figure 4C and E). Meanwhile, GDP does not contribute to this change, but rather shows subtle decrease of the kinetic energy and increase of the bonded energy (Figure 4D).

The nonbonded energy between GDP and P_i_ is mainly attributed to Coulomb energy (see **Figure S4**) and keeps positive value in the time domain (see yellow line in Figure 4E). Retained repulsive interaction between GDP and P_i_ can be explained by a spatial constraint that P_i_ is prohibited to be released from the Ras by GAP-binding and thus is bound around GDP. This observation supports the assumption of the conjecture^12^ that the repulsive Coulomb interaction is the source of mechanical work.

Meanwhile, the Coulomb interaction energy calculated from our atomistic simulations is non-repulsive in total. The non-repulsive, negative part of this energy comes from intra-P_i_ and intra-GDP Coulomb interactions (see orange and pink lines in Figure 4E). Their relaxation also contributes to increases of kinetic energies of GDP and P_i_. Our simulations suggest that the conjecture has room to be improved from a microscopic viewpoint.

Decreasing nonbonded energy partly contributes to increase of bonded energy in the time domain between 0 and 0.01 ps (see Figure 4A). Relaxation of the stored bonded energy starts around 0.02 ps, then bringing about the additional kinetic energy increase, which ranges from 0.02 ps to 0.1 ps. This step was not referred to in the original conjecture and newly found in our atomistic MD simulation, since the conjecture was made from the energy transfer mechanism obtained from analyzing a simple physical model rather than complicated full-atomistic one as in this study.

Finally, each mechanical energy reaches the maximum raise by 0.1 ps and is relaxed to the equilibrium state within several picoseconds (see Figure 4). It is important to note that the kinetic energy gained via the hydrolysis reaction may be partly thermalized and dissipated without doing mechanical work. Kinetic energy of GDP and P_i_ increases from 41 to 64 kcal/mol by the first 0.01 ps (see Figure 4A). This increase is approximately two-fold greater than well-known macroscopic free energy change (ca. 12 kcal/mol) via triphosphate nucleotide hydrolysis and, naturally, than the expected contribution to free energy change estimated in the conjecture^12^, which is at maximum a half of total free energy difference of the hydrolysis reactions.

Recalling the relationship with the conjecture, it should be noted that nonbonded energy acting between GDP and P_i_ is subtly weaken via configurational relaxation but kept positive at the end of the 106-ps NVE simulation. According to one of the two important remarks in the original conjecture^12^, retention of such repulsive Coulomb interaction is the source of mechanical work, which brings about consecutive collisions with then the neighboring chemical groups. However, it is not apparent that the repulsive interaction can do work actually.

This remark implicitly has such an assumption that the kinetic energy of P_i_ and GDP retains the directionality of velocity without thermalization for a certain period. Then, we examined the crucial assumption above by analyzing angular correlation coefficients (ACC) for pairs of atomic velocities. For the GDP system, subsets *A* and *B* are for P_i_ and GDP, respectively (see **Eq. 1** and **Eq. 2** for definition). Four time domains, annotated by *T* in **Eq. 1**, (0-0.1 ps; 0.1-1 ps; 1-6 ps; 6-106 ps) are individually considered. A set of ACC values are illustrated in Figure 5 and Table 2.

**Figure 5.**
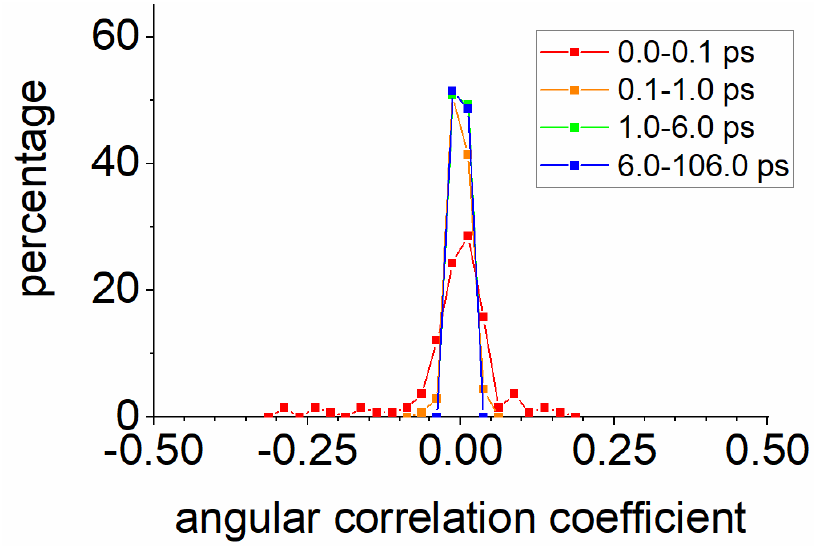
Angular correlation coefficients calculated by using the atomic velocities of atoms in GDP and H_2_PO_4_^−^. The four time domains are distinguished by colors.

**Table 2.** Statistical analyses of angular correlation coefficients of velocities calculated for a pair of atoms from GDP and P_i_.

| time domain | min. | max. | ave. | s.d. |
| --- | --- | --- | --- | --- |
| 0-0.1 ps | −2.9E−01 | 1.6E−01 | −6.3E−03 | 4.4E−03 |
| 0.1-1 ps | −6.2E−02 | 4.2E−02 | −8.1E−04 | 2.1E−04 |
| 1-6 ps | −2.1E−02 | 2.1E−02 | −4.5E−05 | 6.9E−05 |
| 6-106 ps | −2.2E−02 | 2.4E−02 | 9.0E−05 | 7.2E−05 |

Figure 5 clearly indicates rapid loss of relative directionality of velocity associated with kinetic energy within several picoseconds. We can find relatively large ACC values whose magnitudes are greater than 0.1 in the time domain of 0-0.1 ps. However, such values are absent in the following time domains. Actually, as shown in Table 2, the magnitude of ACC is lowered to order of 10^−2^ by the first 1 ps. Therefore, the kinetic energy of P_i_ and GDP cannot retain the velocity directionality by the timing to do work for biologically important processes which progress on the timescale of several micro seconds or longer, but rather promptly become randomized heats at atomic level. Then we could suppose that the GTP hydrolysis does not do work on the Ras-GAP system by bringing about consecutive collisions of GDP and P_i_ with the neighboring chemical groups (see also the subsections below).

### GTP-GDP conversion increases potential energy of P-loop in Ras protein

We next focus on functional regions of Ras protein (switch I, switch II, P-loop and α-helix 3, shown in Figure 1B) close to the GTP hydrolysis site, and analyze the effects of GTP-GDP conversion on these regions.

Among the four functional regions, only the P-loop shows significant changes through the reaction (see **Figures S5**, **S6** and **S7** in Supporting Information for similar analyses for Switch I, Switch II and α-helix 3, respectively). The kinetic and bonded interaction energies of P-loop rapidly return to their initial level of the values within several picoseconds (Figure 6A and B), while the nonbonded interaction energy of the P-loop starts to increase at around 0.01 ps and retains the change even after the relaxation of other two energy terms (Figure 6C).

**Figure 6.**
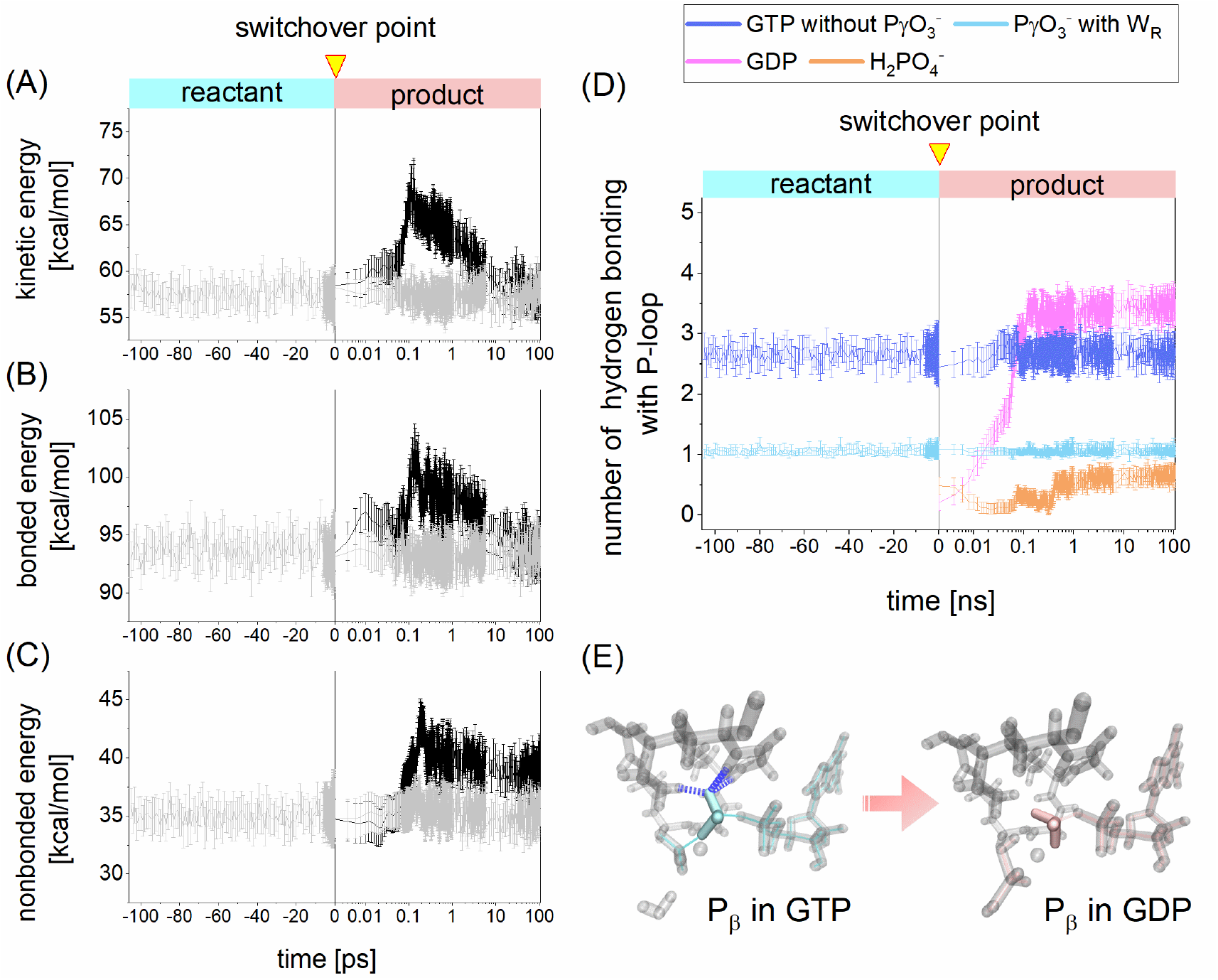
Energetic and structural changes of P-loop in Ras through GTP-GDP conversion. (A)-(C) temporal changes of kinetic, bonded and nonbonded energies, where Ras-GTP-GAP system and Ras-GDP-GAP system are annotated by grey and black lines, respectively. (D) hydrogen bond formation between P-loop and reactant (GTP with reactant water molecule (W_R_))/product (GDP and inorganic phosphate (P_i_)). Ras-GTP-GAP system is changed into Ras-GDP-GAP system at the switchover point, 0 ps on the time course. (E) Representative illustration for configurational change of P_β_ via conversion from GTP and GDP. An error bar indicates 95% confidential interval.

Here the effect of GTP hydrolysis reaction on the Ras protein appears as the increase of interaction energy of the P-loop, thus denoting that mechanical work generated via the GTP hydrolysis converts into the local energy in the P-loop and is stored inside the Ras protein. It has been known that the P-loop of Ras protein is involved in the GTP hydrolysis process by redistributing the charge density of GTP.^34^ Meanwhile, we newly found the alternative role of the P-loop in the post-GTP hydrolysis process as the storage of mechanical energy. Thus, we will focus on energetic and structural properties of the P-loop in the following.

The energy storage of P-loop may be compensated by the energy release of GDP and P_i_. The value of nonbonded energy for the P-loop increases by 5.3 kcal/mol through the back 106-ps NVE simulation, namely, b-NVE (34.9 ± 1.7 kcal/mol at 0 ps; 40.2 ± 1.4 kcal/mol at 106 ps). This increase of the nonbonded energy may be obtained from the decrease in the potential energy of GPD plus P_i_ (sum of the bonded and nonbonded energies shown in Figure 4A). The value of the potential energy decreases by 88.3 kcal/mol at 106 ps via the b-NVE. Meanwhile, the total potential energy of the Ras-GDP-GAP system decrease by 83.1 kcal/mol at 106 ps via the b-NVE (Figure 7B): this change is mostly offset by the increase of total kinetic energy of 82.4 kcal/mol (Figure 7A and 7C) due to nature of the NVE simulation. The remaining 5 kcal/mol seems to be stored as nonbonded energy of P-loop.

**Figure 7.**
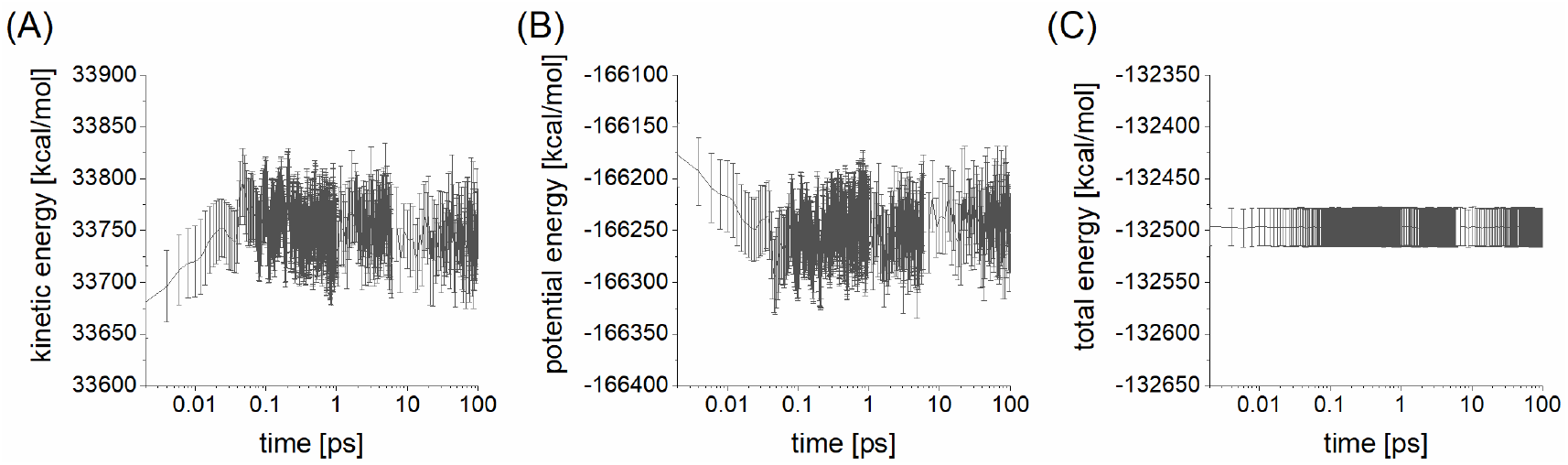
Mechanical energy of Ras-GDP-GAP system. (A) kinetic energy. (B) potential energy (bonded energy plus nonbonded one). (C) total energy (sum of kinetic and potential energies). An error bar indicates 95% confidential interval.

Here, the 5 kcal/mol is about a half of the free energy change via the hydrolysis reaction under the physiological condition (ca. 12 kcal/mol). Although there would be additional physicochemical processes which contribute to the free energy change, they are not considered in the framework of our simulations. In the realistic system, the hydrolysis reaction proceeds in a finite time duration accompanied by structural and energetic relaxations of the reaction intermediates. Meanwhile, we instantaneously simulated the GTP hydrolysis by the SF2MD because we aimed to examine Ross’ conjecture concerning the roles of repulsive Coulomb interactions between GDP and P_i_, which result from the hydrolysis reaction.

It is interesting to find the remaining portion of the total free energy change within the whole reaction processes, whereas, as remarked in the Introduction of this manuscript, it is not feasible to straightforwardly examine the *ab initio* molecular dynamics of the hydrolysis reaction with the computational resources of today. Thus, we would like to consider the contribution of such relaxation processes in the future.

The GTP hydrolysis leads to rearrangement of intermolecular hydrogen bond (HB) formation between the P-loop and GDP with P_i_. Figure 6D represents temporal change of HB formation between the P-loop and GTP with W_R_, and that between the P-loop and GDP with P_i_. GDP and P_i_ are separately examined in analyses of HB formation with the P-loop. As the counterparts in Ras-GTP-GAP system, GTP without P_γ_O_3_^−^ and P_γ_O_3_^−^ with W_R_ were considered in similar HB formation analyses. P_i_ forms smaller number of HB with P-loop than P_γ_O_3_^−^ with W_R_. Meanwhile, we can find that GDP finally forms greater number of HB with P-loop by ca. 0.5 than GTP without P_γ_O_3_^−^.

The number of HBs between P-loop and GDP shows significant decrease at the force field switchover point. This is due to configurational change of P_β_O_3_^−^ through modeling the product states (see Figure 6E for a representative configurations of P_β_O_3_^−^). We can find that P_β_O_3_^−^ at the point is reoriented to break hydrogen bonding with P-loop.

The HB rearrangement discussed above is the structural hallmark of the energetic change of the P-loop via the GTP-GDP conversion. The role of the P-loop could be experimentally tested by examining the structural difference in the P-loop associated HB formation between the GTP-bound Ras-GAP system and GDP-bound one. Recalling the previous study on the GTPase mechanism of Ras^34^, we suppose that the time-resolved infrared difference spectroscopy methods can be a feasible approach for the purpose^35^.

It is a straightforward expectation that rearrangement of HBs between P-loop and GDP or P_i_ can be used for the source of mechanical work, instead of a pair of GDP and P_i_ discussed with regard to Figure 4. From this point of view, we reconsider the remark for consecutive collisions discussed above, again. We then analyzed angular correlation coefficients (ACC) of velocities for pairs of atoms in P-loop and GDP and for those in the P-loop and P_i_ (Figure 9A and B). Relatively large ACC values are found in the time domain of 0-0.1 ps, although such values are absent in the following time domains as in the case of ACC for GDP-P_i_ (Figure 5). Actually, as shown in Table 3, the magnitude of ACC is lowered by order to 10^−2^ by the first 1 ps.

**Figure 8.**
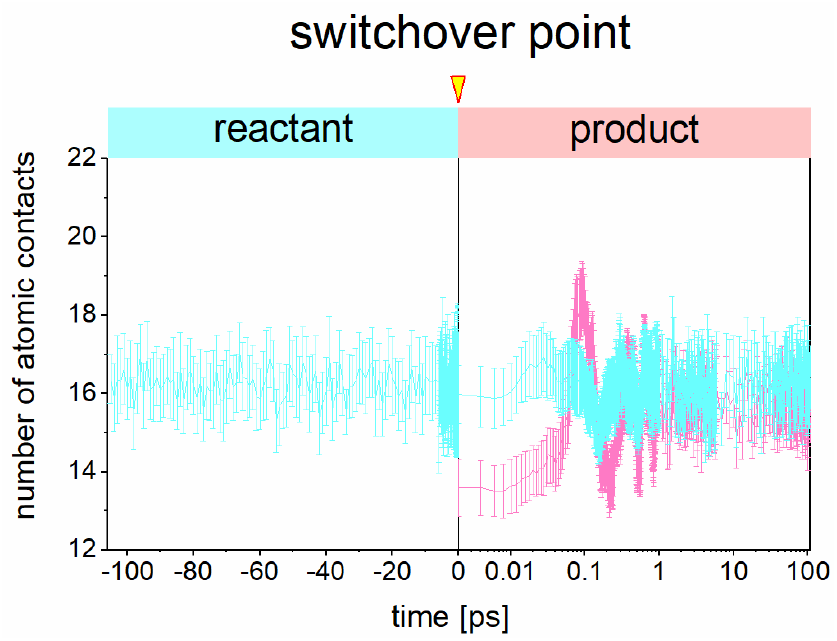
Temporal change of atomic contacts between P-loop and GDP plus P_i_ through GTP-GDP conversion. GTP in Ras-GTP-GAP system and GDP plus P_i_ in Ras-GDP-GAP system are distinguished by blue and pink lines, respectively. An error bar indicates 95% confidential interval.

**Figure 9.**
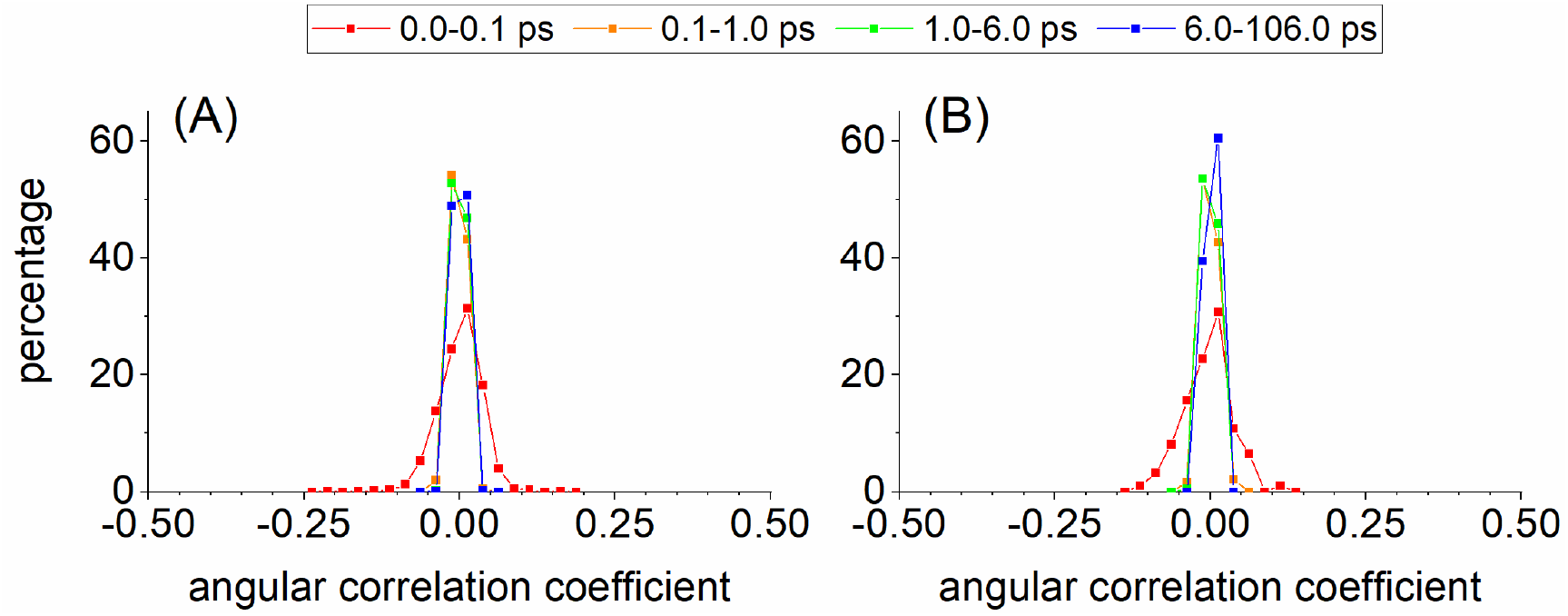
Angular correlation coefficients of velocities calculated between (A) atoms in P-loop and GDP, and (B) atoms in the P-loop and P_i_. The four time domains are distinguished by colors.

**Table 3.** Statistical analyses of angular correlation coefficients of velocities calculated for a pair of atoms from P-loop and GDP, and for a pair of atoms from the P-loop and P_i_.

| time domain | P-loop and GDP |  |  |  |
| --- | --- | --- | --- | --- |
|  | min. | max. | ave. | s.d. |
| 0-0.1 ps | −2.2E−01 | 1.6E−01 | 5.3E−04 | 1.2E−03 |
| 0.1-1 ps | −4.3E−02 | 3.1E−01 | −1.7E−03 | 1.1E−04 |
| 1-6 ps | −3.0E−02 | 2.6E−02 | −4.6E−04 | 6.6E−05 |
| 6-106 ps | −3.0E−02 | 2.7E−02 | 4.7E−05 | 6.9E−05 |

| time domain | P-loop and H<sub>2</sub>PO<sub>4</sub><sup>−</sup> |  |  |  |
| --- | --- | --- | --- | --- |
|  | min. | max. | ave. | s.d. |
| 0-0.1 ps | −1.1E−01 | 1.2E−01 | −5.3E−03 | 1.5E−03 |
| 0.1-1 ps | −3.0E−02 | 3.7E−02 | −8.2E−04 | 1.2E−04 |
| 1-6 ps | −2.5E−02 | 2.4E−02 | −3.2E−04 | 7.0E−05 |
| 6-106 ps | −2.3E−02 | 2.2E−02 | 1.5E−03 | 5.7E−05 |

Considering the loss of the velocity directionality associated with kinetic energy, we concluded that kinetic energy of P-loop and GDP plus P_i_ cannot be a source of mechanical work. Similar analyses were performed over all residues of Ras and GAP proteins, while we could not find significant effects of the hydrolysis reaction (see **S4** in Supporting Information). Thus, the remark for generation mechanism of mechanical work via consecutive collisions with neighboring chemical groups does not seem to be essential for understanding of the role of GTP hydrolyses in the Ras-GAP system.

The energetic and structural changes discussed above are sustained even in the following 20-ns unbiased NVE MD simulations (Figure 10). It could be expected that the GTP-GDP conversion affects following reaction steps such as Ras-GAP dissociation and/or P_i_ release from Ras. Considering these points is interesting to clarify mechanisms of re-activation process of Ras, which begins with GAP dissociation. However, the subject of the present study is to examine whether repulsive Coulomb interaction acting between GDP and P_i_ does work to the system. Further investigation for this question is beyond the scope of this study, thus being left for future studies.

**Figure 10.**
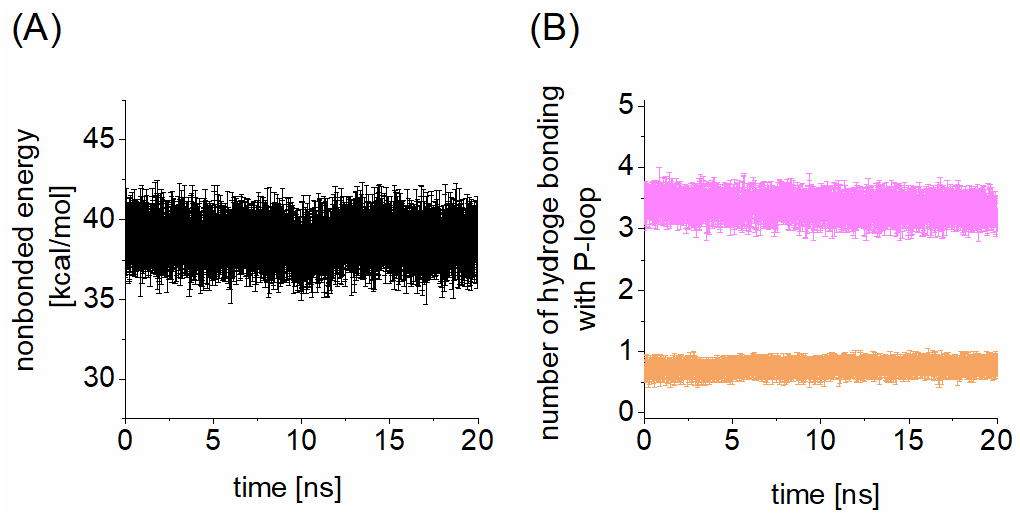
Energetic and structural analyses for extended 20-ns NVE MD simulations for Ras-GDP-GAP system. (A) nonbonded energy of P-loop. (B) hydrogen bond formation between P-loop and products (GDP and P_i_, shown by orange and pink lines, respectively). An error bar indicates 95% confidential interval.

### The change associated with the P-loop is brought about via GTP-GDP conversion with no regard to raise of kinetic energy

The transient raise of kinetic energy of the P-loop in these simulations appears to be induced by the impulsive collisions between the P-loop and GDP with P_i_. The kinetic energy of the P-loop shows immediate increases within 1 ps (Figure 6A). This change apparently follows the raise of kinetic energy of GDP and P_i_, which occurs by the first 0.1 ps (compare Figure 4C and Figure 6A) and is simultaneous with impulsive increase of the number of atomic contacts between the P-loop and GDP plus P_i_ (Figure 8).

It is noted that the kinetic energy of GDP and P_i_ is mainly transferred to the P-loop through local interactions between individual atoms. The Ross conjecture^12^ describes GDP and P_i_ as rigid bodies so that the transfer of their kinetic energies is conceptually associated with the translational motions of their centers of mass. However, the raises of kinetic energy of the GDP and P_i_ mainly come from the individual atomic motions relative to their centers of mass (**Figure S8**), denoting that the rigid-body translational motions are unimportant for the energy transfer between P-loop and GDP with P_i_.

Although the energy and structure analyses above challenges the essential assumption of the conjecture^12^, they still support another important remark in the conjecture. The impulsive collisions with neighboring chemical groups might do work for the increase of nonbonded energy of the P-loop, whereas a causal relationship between the two microscopic events is elusive so far. Then, we examine such a relationship by quenching the increase of kinetic energy with the two different simulation procedures.

First, the kinetic energy was absorbed into the thermostat during the initial 1-ps in b-NVE (Figure 11A and B). The bonded and nonbonded energies of P-loop start to increase around 0.01 ps and only the latter retains the change at the end of the b-NVE simulations (Figure 11C and D). The rearrangement of the HB formation was observed even under this heat-absorbing simulation condition (Figure 11E). It is noted that completion of the rearrangement takes 1 ps (compare Figure 11D with Figure 6C). Suppressing increase of kinetic energy of GDP with P_i_ may delay their encounter with the P-loop and thus retards the rearrangement of HB formation by tenfold. Nonetheless, the HB rearrangement is a fast process which finishes by the first 1 ps, even under this simulation condition. This denotes that the rearrangement of HB formation proceeds with no regard to increase of kinetic energy via GTP-GDP conversion and is driven by the energetic relaxation on the reshaped potential energy surface.

**Figure 11.**
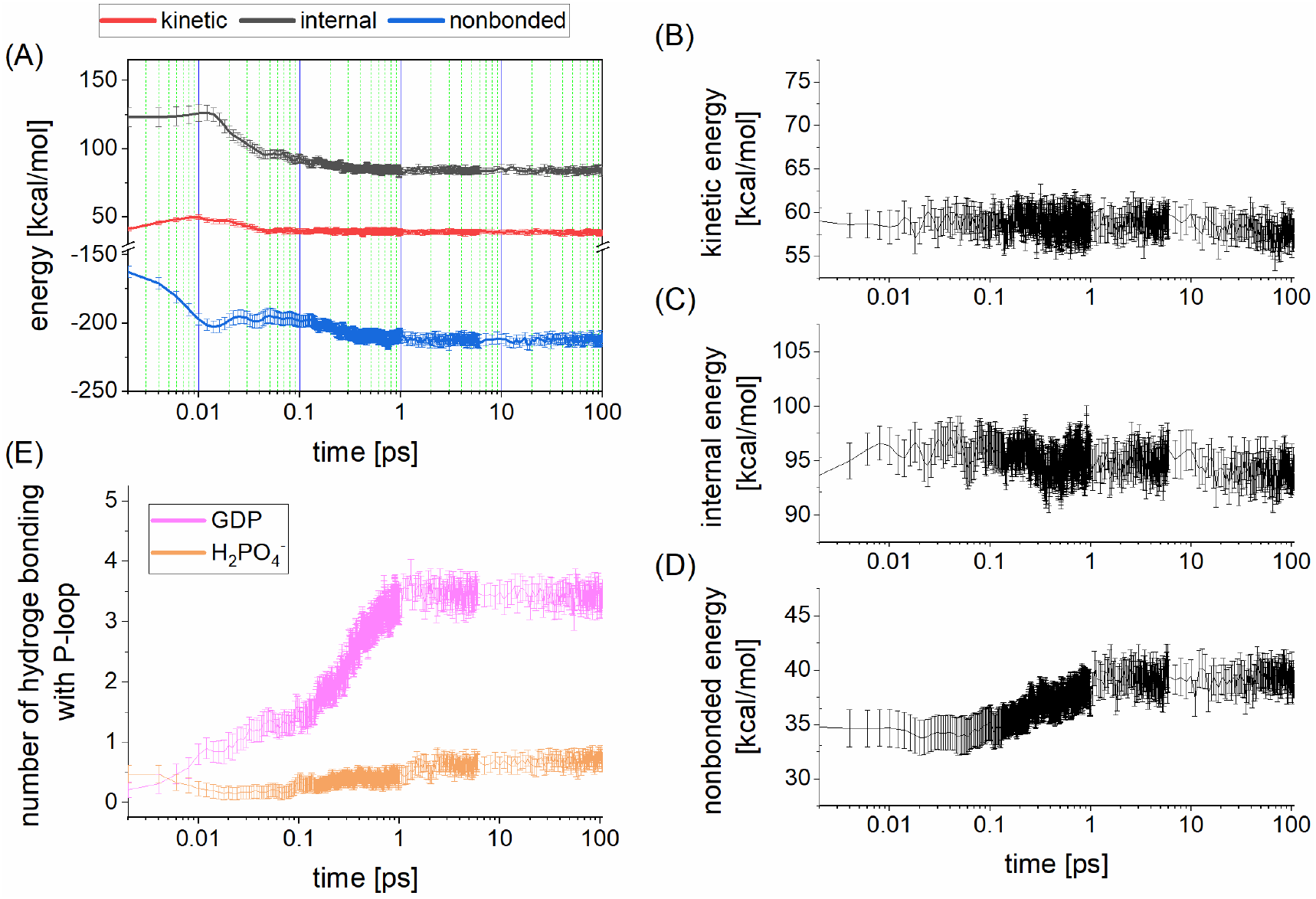
Energetic and structural changes of Ras-GDP-GAP system with heat absorption by interacting with the thermal bath during the first 1 ps. (A) mechanical energy of GDP plus P_i_. (B)-(D) kinetic, bonded and nonbonded energies of the P-loop. (E) hydrogen bond formation between P-loop and products (GDP and P_i_). An error bar indicates 95% confidential interval.

Secondly, P_i_ and GDP are energetically optimized in advance of the b-NVE, aiming to relax the repulsive interaction which can be converted into kinetic energy (Figure 12). We find that this simulation suppresses increase of kinetic energies for GDP with P_i_ and P-loop (Figure 12A and B). Bonded energy of P-loop appears to be nearly constant in the b-NVE simulations, while the nonbonded energy of P-loop starts to increase at around 0.1 ps and retains the change at the end of the NVE simulations (Figure 12C and D). The rearrangement of the HB formation was completed by 1 ps (Figure 12E). We find that the number of HBs between GDP and P-loop starts from ca. 3, a relatively large value compared to the cases of the unbiased b-NVE discussed above and the kinetic energy absorption b-NVE discussed just above (compare Figure 12E with Figure 6D and Figure 11E, respectively). Such a preformed hydrogen bonding should be due to the preliminary energetic optimization. It can be said that the rearrangement of HB is a thermodynamically spontaneous process on potential energy surface of Ras-GDP-GAP system.

**Figure 12.**
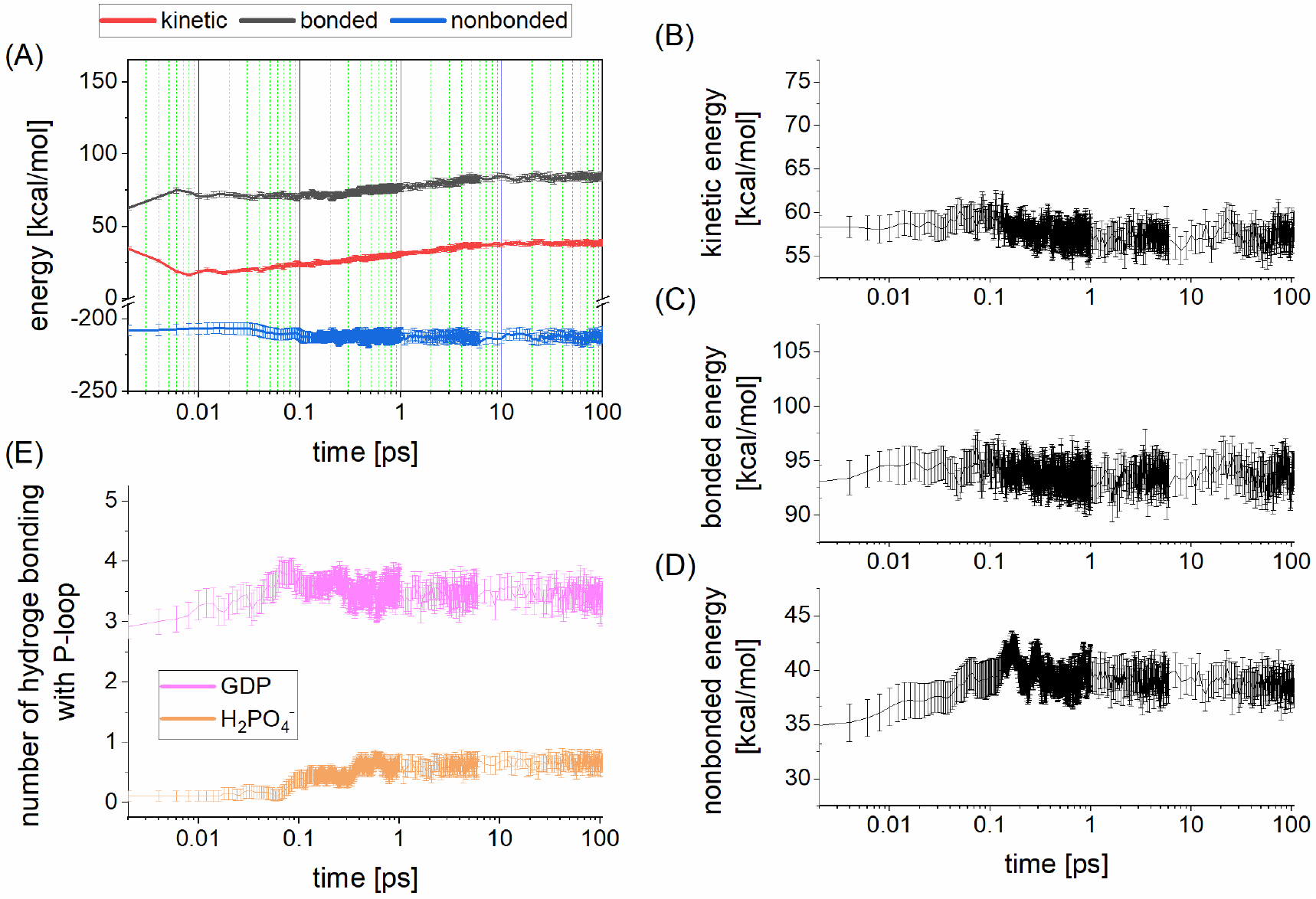
Energetic and structural changes of Ras-GDP-GAP system after preliminary optimization of GDP-P_i_ configuration. (A) mechanical energy of GDP plus P_i_. (B)-(D) kinetic, bonded and nonbonded energies of the P-loop. (E) hydrogen bond formation between P-loop and products (GDP and P_i_). An error bar indicates 95% confidential interval.

These two additional simulations provide the observation that 5 kcal/mol energy storage in P-loop is brought about via GTP-GDP conversion, without regard to transfer of kinetic energy of GDP and P_i_ to P-loop. Therefore, the earlier remark for generation mechanism of mechanical work via impulsive collisions with neighboring chemical groups is also challenged with regard to understanding of the role of GTP hydrolyses in the Ras-GAP system.

Triphosphate nucleotide hydrolysis is an exothermic reaction so that, in the earlier MD simulation study^36^, the effect of GTP hydrolysis on Ras protein was represented by randomly adding kinetic energy to GDP. An assumption that underlies such a physicochemical simulation may be that a substantial amount of kinetic energy is transferred to a specific reaction coordinate^37, 38^ and drives conformational changes via vibrational motion along the reaction coordinate or mode.

Meanwhile, judging from the present results illustrated above, it is unlikely that such an added kinetic energy can be kept on a specific reaction coordinate and do work for slower biochemical processes in Ras-GAP system. Our SF2MD simulations show rapid dissipation of increased kinetic energy and prompt loss of directionality of kinetic momenta within several picoseconds. Whereas the biochemical processes which follow the GTP hydrolysis, such as GAP dissociation and P_i_ release from Ras, progress on the timescale of several micro seconds or longer, thus being 10^6^-fold slower than the raise of kinetic energy via GTP-GDP conversion and the thermalization.

Recalling these observations, we could suppose that ATPase/GTPase proteins are not a single-molecular transducer which straightforwardly converts kinetic energy obtained from ATP/GTP hydrolysis reaction into mechanical work. This claim may be supported by the experimental observations achieved from the study on F_1_-ATPase^39^, where the authors consider that the functional role of the ATP hydrolysis for mechanical force generation is promotion of inorganic phosphate release, and the ATP hydrolysis as exothermic reaction is not main driving force for the functional expression.

From microscopic viewpoints, the atomistic simulation study discussed that such a mechanical work is generated by relaxation of structural distortion of F_1_-ATPase, which accompanies the release of inorganic phosphate from one of the six subunits^40^. By recalling that Ras takes the inactive conformation by binding to GDP in the absence of P_i_,^41^ we could suppose that the release of P_i_ from Ras brings about such a conformational change. This speculation will be considered in the future by taking into account the energy increase of 5 kcal/mol in the P-loop, namely, phosphate binding loop.

Then, the observations obtained in the present study highlight such an activation mechanism via triphosphate nucleotide hydrolysis that the chemical conversion of GTP/ATP into GDP/ADP itself plays an important role to do work for a system. Effective change of the potential energy surface of the molecular system brings about the emergent steric interactions between the product molecules and the neighboring chemical groups in proteins. Spontaneously relaxing such an interaction toward thermodynamically stable configurations does work to express the molecular function. It seems quite rational for physicochemical designs of biomacromolecules that their biochemical functions can be rigorously regulated by changing relative thermodynamic stability among configuration repertoire of the system, virtually independently of way of directionless and dissipative kinetic energy gain.

## Conclusion

In the present study, we aimed to obtain deeper physicochemical insights into the generation mechanism of mechanical work via triphosphate nucleotide hydrolysis by considering one of GTPase proteins, Rat sarcoma (Ras). The earlier conjecture focusing on roles of repulsive Coulomb interaction acting between GDP and inorganic phosphate^12^, P_i_, as a source of mechanical work was examined as a working hypothesis.

First, the conventional switching force field method was technically improved to study complicated chemical reactions and applied to simulating non-equilibrium energy relaxation process driven by GTP hydrolysis reaction of Ras. Within period of several picoseconds, we observed rapid conversion of Coulomb interaction energy acting between GDP and P_i_ into their kinetic energy and also increase of nonbonded energy of P-loop in Ras protein by 5 kcal/mol, which was retained by the following 20 ns.

By testing the two essential remarks given in the earlier conjecture, we found that neither of them are associated with generation mechanism of mechanical work via the GTP-GDP conversion. Instead, our analyses indicate that GTP-GDP conversion itself straightforwardly does work in the form of potential energy storage in P-loop of Ras protein and may cause approximately half of the overall free energy changes via the GTP hydrolysis reaction.

It has been known^34^ that the P-loop in the Ras protein promotes the GTP hydrolysis by redistributing the charge density of GTP. Meanwhile, the present study newly showed the alternative role of the P-loop in the post-GTP hydrolysis process, then implying that the P-loop is involved in the overall process of functional regulation of the Ras protein. Besides, we could suppose that similar observations would be obtained for other ATPase and GTPase because the P-loop motif is commonly found in various ATPase and GTPase proteins^42^.

The observations that we obtained evokes that the work generation mechanism of ATPase and GTPase is imprinted in the repertoire of their atomic structures and mostly independent of microscopic processes of atomic heat generation by ATP/GTP hydrolysis reactions. One of the important roles of the triphosphate nucleotide hydrolysis in mechanical work generation may be the production of emergent steric interactions that do work via the relaxation, namely, a shift of biomolecular system to non-equilibrium state by reshaping the potential energy landscape.

## Acknowledgements

This work was supported by a Grant-Aid for Scientific Research on Innovative Areas “Chemistry for Multimolecular Crowding Biosystems” (JSPS KAKENHI Grand No. JP17H06353). Parts of MD simulations discussed here were performed by using supercomputers at the Research Center for Computational Science, Okazaki Research Facilities, National Institutes of Natural Sciences, Japan. Partial support by MEXT Quantum Leap Flagship Program (MEXT QLEAP; Grant Number JPMXS0120330644) is also acknowledged.

## References

1. J. Howard, Nature, 1997, 389, 561–567.

2. E. Nogales and H. W. Wang, Curr. Opin. Struc. Biol., 2006, 16, 221–229.

3. E. Gouaux and R. MacKinnon, Science, 2005, 310, 1461–1465.

4. B. S. Strunk and K. Karbstein, RNA, 2009, 15, 2083–2104.

5. Y. Takai, T. Sasaki and T. Matozaki, Physiol. Rev., 2001, 81, 153–208.

6. K. L. Pierce, R. T. Premont and R. J. Lefkowitz, Nat. Rev. Mol. Cell Bio., 2002, 3, 639–650.

7. P. Nogly, T. Weinert, D. James, S. Carbajos, D. Ozerov, I. Schapiro, G. Schertler, R. Neutze and J. Standfuss, Acta Crystallogr. A, 2018, 74, E170–E170.

8. N. Shibayama, A. Sato-Tomita, M. Ohki, K. Ichiyanagi and S. Y. Park, P. Natl. Acad. Sci. USA., 2020, 117, 4741–4748.

9. R. Salomon-Ferrer, A. W. Gotz, D. Poole, S. Le Grand and R. C. Walker, J. Chem. Theory Comput., 2013, 9, 3878–3888.

10. D. K. Simanshu, D. V. Nissley and F. McCormick, Cell, 2017, 170, 17–33.

11. T. Schweins, M. Geyer, K. Scheffzek, A. Warshel, H. R. Kalbitzer and A. Wittinghofer, Nat. Struct. Biol., 1995, 2, 36–44.

12. J. Ross, J. Phys. Chem. B, 2006, 110, 6987–6990.

13. M. Takayanagi and M. Nagaoka, Theor. Chem. Acc., 2011, 130, 1115–1129.

14. M. Takayanagi, H. Okumura and M. Nagaoka, J. Phys. Chem. B, 2007, 111, 864–869.

15. J. A. Maier, C. Martinez, K. Kasavajhala, L. Wickstrom, K. E. Hauser and C. Simmerling, J. Chem. Theory Comput., 2015, 11, 3696–3713.

16. P. G. Kusalik and I. M. Svishchev, Science, 1994, 265, 1219–1221.

17. W. L. Jorgensen, J. Chandrasekhar, J. D. Madura, R. W. Impey and M. L. Klein, J. Chem. Phys., 1983, 79, 926–935.

18. I. S. Joung and T. E. Cheatham, III, J. Phys. Chem.y B, 2009, 113, 13279–13290.

19. I. S. Joung and T. E. Cheatham, J. Phys. Chem. B, 2008, 112, 9020–9041.

20. K. L. Meagher, L. T. Redman and H. A. Carlson, J Comput. Chem., 2003, 24, 1016–1025.

21. O. Allner, L. Nilsson and A. Villa, J. Chem. Theory Comput., 2012, 8, 1493–1502.

22. M. G. Khrenova, B. L. Grigorenko, A. B. Kolomeisky and A. V. Nemukhin, J. Phys. Chem. B, 2015, 119, 12838–12845.

23. M. J. Frisch, G. W. Trucks, H. B. Schlegel, G. E. Scuseria, M. A. Robb, J. R. Cheeseman, G. Scalmani, V. Barone, G. A. Petersson, H. Nakatsuji, X. Li, M. Caricato, A. V. Marenich, J. Bloino, B. G. Janesko, R. Gomperts, B. Mennucci, H. P. Hratchian, J. V. Ortiz, A. F. Izmaylov, J. L. Sonnenberg, Williams, F. Ding, F. Lipparini, F. Egidi, J. Goings, B. Peng, A. Petrone, T. Henderson, D. Ranasinghe, V. G. Zakrzewski, J. Gao, N. Rega, G. Zheng, W. Liang, M. Hada, M. Ehara, K. Toyota, R. Fukuda, J. Hasegawa, M. Ishida, T. Nakajima, Y. Honda, O. Kitao, H. Nakai, T. Vreven, K. Throssell, J. A. Montgomery Jr., J. E. Peralta, F. Ogliaro, M. J. Bearpark, J. J. Heyd, E. N. Brothers, K. N. Kudin, V. N. Staroverov, T. A. Keith, R. Kobayashi, J. Normand, K. Raghavachari, A. P. Rendell, J. C. Burant, S. S. Iyengar, J. Tomasi, M. Cossi, J. M. Millam, M. Klene, C. Adamo, R. Cammi, J. W. Ochterski, R. L. Martin, K. Morokuma, O. Farkas, J. B. Foresman and D. J. Fox, Gaussian 09 Rev. A.02, Wallingford, CT, 2016.

24. J. M. Wang, R. M. Wolf, J. W. Caldwell, P. A. Kollman and D. A. Case, J. Comput. Chem., 2004, 25, 1157–1174.

25. K. Scheffzek, M. R. Ahmadian, W. Kabsch, L. Wiesmuller, A. Lautwein, F. Schmitz and A. Wittinghofer, Science, 1997, 277, 333–338.

26. H. J. C. Berendsen, J. P. M. Postma, W. F. Vangunsteren, A. Dinola and J. R. Haak, J. Chem. Phys., 1984, 81, 3684–3690.

27. D. A. Case, D. S. Cerutti, T. E. Cheatham, III, T. A. Darden, R. E. Duke, T. J. Giese, H. Gohlke, A. W. Goetz, D. Greene, N. Homeyer, S. Izadi, A. Kovalenko, T. S. Lee, S. LeGrand, P. Li, C. Lin, J. Liu, T. Luchko, R. Luo, D. Mermelstein, K. M. Merz, G. Monard, H. Nguyen, I. Omelyan, A. Onufriev, F. Pan, R. Qi, D. R. Roe, A. Roitberg, C. Sagui, C. Simmerling, W. M. Botello-Smith, J. Swails, R. C. Walker, J. Wang, R. M. Wolf, X. Wu, L. Xiao, D. M. York and P. A. Kollman, Amber 17, University of California, San Francisco, 2017.

28. S. Le Grand, A. W. Gotz and R. C. Walker, Comput. Phys. Commun., 2013, 184, 374–380.

29. . D. A. Case, K. Belfon, I. Y. Ben-Shalom, S. R. Brozell, D. S. Cerutti, T. E. Cheatham, III, V. W. D. Cruzeiro, T. A. Darden, R. E. Duke, G. Giambasu, M. K. Gilson, H. Gohlke, A. W. Goetz, R. Harris, S. Izadi, S. A. Izmailov, K. Kasavajhala, A. Kovalenko, R. Krasny, T. Kurtzman, T. S. Lee, S. LeGrand, P. Li, C. Lin, J. Liu, T. Luchko, R. Luo, V. Man, K. M. Merz, Y. MIao, O. Mikhailovskii, G. Monard, H. Nguyen, A. Onufriev, F. Pan, S. Pantano, R. Qi, D. R. Roe, A. Roitberg, C. Sagui, S. Schott-Verdugo, J. Shen, C. Simmerling, N. R. Skrynnikov, J. Smith, J. Swails, R. C. Walker, J. Wang, L. Willson, R. M. Wolf, X. Wu, Y. Xiong, Y. Xue, D. M. York and P. A. Kollman, Amber 20, University of California, San Francisco, 2020.

30. W. Thiel, Wires. Comput. Mol. Sci., 2014, 4, 145–157.

31. R. C. Walker, M. F. Crowley and D. A. Case, J. Comput. Chem., 2008, 29, 1019–1031.

32. K. Nam, J. L. Gao and D. M. York, J. Chem. Theory Comput., 2005, 1, 2–13.

33. W. Humphrey, A. Dalke and K. Schulten, J. Mol. Graphics Model., 1996, 14, 33–38.

34. C. Allin and K. Gerwert, Biochemistry, 2001, 40, 3037–3046.

35. V. A. Lorenz-Fonfria, Chem. Rev., 2020, 120, 3466–3576.

36. K. Ogata, J. W. Shen, S. Sugawa and S. Nakamura, B. Chem. Soc. Jpn., 2012, 85, 1318–1328.

37. K. Moritsugu, O. Miyashita and A. Kidera, Phys. Rev. Lett., 2000, 85, 3970–3973.

38. S. Tanaka, J. Phys. Soc. Jpn., 2012, 81.

39. C. B. Li, H. Ueno, R. Watanabe, H. Noji and T. Komatsuzaki, Nat. Commun., 2015, 6.

40. K. Okazaki and G. Hummer, P. Natl. Acad. Sci. USA, 2013, 110, 16468–16473.

41. S. Matsumoto, N. Miyano, S. Baba, J. L. Liao, T. Kawamura, C. Tsuda, A. Takeda, M. Yamamoto, T. Kumasaka, T. Kataoka and F. Shima, Sci. Rep-Uk, 2016, 6.

42. M. Saraste, P. R. Sibbald and A. Wittinghofer, Trends Biochem. Sci., 1990, 15, 430–434.

