## Supporting Information for "Elucidating Microscopic Events Driven by GTP Hydrolysis Reaction in Ras-GAP System with Semi-reactive Molecular Dynamics Simulation: Alternative Role of Phosphate Binding Loop as Mechanical Energy Storage"

*Ikuo Kurisaki

*Shigenori Tanaka

#### SI 1. System setup

For the Ras-GTP-GAP system, the atomic coordinates were energetically relaxed by the following molecular mechanics (MM). First, steric clashes in the system were removed by the three-step MM simulation. The initial MM simulation was for $Mg^{2+}$ and GTP. The next one was for all hydrogen atoms in proteins and ligands. The last one was for all atoms in the system. Each MM simulation consists of 1500 steps of steepest descent method followed by 48500 steps of conjugate gradient method. Atoms except for the $Mg^{2+}$ and GTP and those except for all hydrogen atoms in proteins and ligands were restrained by harmonic potential with force constant of 100 kcal/mol around the initial atomic coordinates of the simulation during the first and second MM simulations, respectively.

#### SI 2. MD simulations for structure relaxation

The temperature and density of the Ras-GTP-GAP system were relaxed through the following five MD simulations: NVT (0.001 to 1 K, 0.1 ps) → NVT (1 K, 0.1 ps) → NVT (1 to 300 K, 20 ps) → NVT (300 K, 20 ps) → NPT (300 K, 300 ps, 1 bar). In each of the five MD simulations, the atomic coordinates except for water and $Na^+$ molecules were restrained by harmonic potential with force constant of 100 [kcal/mol/Å$^2$] around the initial atomic coordinates of the simulation.

The first two NVT MD simulations and the other MD simulations were performed using 0.01 fs and 2 fs for the time step of integration, respectively. The first two NVT simulations and the following three ones were performed using Berendsen thermostat[1] with a 0.001-ps coupling constant and Langevin thermostat with 1-$ps^{-1}$ collision coefficient, respectively. In the first, second and third NVT simulations, the reference temperature was linearly increased along the time course. The NPT simulation was performed using Langevin thermostat with 1-$ps^{-1}$ of collision coefficient and, Berendsen barostat[1] with 2-ps coupling constant. A set of initial atomic velocities was randomly assigned from the Maxwellian distribution at 0.001 K at the beginning of the first NVT simulation.

Using a set of the atomic coordinates derived from the above relaxation simulation, an Ras-GTP-GAP complex was structurally relaxed in aqueous solution through the following 7-step MD simulations: NVT (0.001 to 1 K, 0.1 ps, 10 kcal/mol/Å) → NVT (1 to 300 K, 0.1 ps, 10 kcal/mol/Å) → NVT (300 K, 10 ps, 10 kcal/mol/Å) → NVT (300 K, 40 ps, 5 kcal/mol/Å) → NVT (300 K, 40 ps, 1 kcal/mol/Å) → NVT (300 K, 40 ps) → NPT (300 K, 1 bar, 20 ns). The last 20-ns NPT simulation was used for the following analyses.

The first two NVT MD simulations and the other MD simulations were performed using 0.01 fs and 2 fs for the time step of integration, respectively. In the first two NVT simulations, the reference temperature was linearly increased along the time-course. In the first 4 steps, Ras, GAP, GTP and $Mg^{2+}$ were restrained by harmonic potentials, whose force constants are shown in the above parentheses, around the initial atomic coordinates. In each NVT simulation, temperature was regulated using Langevin thermostat with 1-$ps^{-1}$ collision coefficient. In the last 20-ns NPT simulation, temperature and pressure were regulated by Berendsen thermostat[1] with a 5-ps coupling constant, and Berendsen barostat[1] with a 0.1-ps coupling constant, respectively. The initial atomic velocities were randomly assigned from the Maxwellian distribution at 0.001 K. The initial atomic velocities were randomly assigned from a Maxwellian distribution at 0.001 [K]. MD trajectories were recorded every 10-ps interval for the following analyses, RMSd calculations.

#### SI 3. Technical advantage of our SF2MD approach in study on ATP and GTP hydrolysis reactions

Triphosphate nucleotide hydrolysis is an exothermic reaction so that, in the earlier MD simulation study[2], the effect of GTP hydrolysis on proteins is practically described by randomly adding kinetic energy to GDP as found. However, this kind of simulation procedure usually raises concerns for reproducibility of physicochemical observations obtained from MD simulations.

Meanwhile, our SF2MD simulation under NVE condition can avoid any randomness upon addition of kinetic energy to GDP and $P_i$ and associated uncertainty: we can definitely simulate atomistic exothermic process via the chemical conversion, which simply depends on atomic coordinates with atomic velocities of the molecular system. Simultaneously, this simulation scheme is designed to significantly suppress thermal noise in atomistic dynamics. We could find essential change triggered by hydrolysis reactions with fewer number of molecular dynamics simulation trajectories.

Furthermore, employing QM/MM simulations for GTP-GDP conversion is useful to perform physicochemically reasonable simulations. We can show it by using MM-based modeling of GDP systems as reference. GTP-GDP conversion was carried out under the MM potential function of Ras-GTP-GAP system, where the QM/MM setup for reactive atoms was turned off. This procedure was applied to the same 49 Ras-GTP-GAP snapshot structures. RMSd values for $P_i$ molecules generated by the MM simulations have RMSd value of $0.64 \pm 0.1$ [Å]. Recalling that those for $P_i$ molecules generated via QM/MM simulations have RMSd value of $0.21 \pm 0.02$ [Å], thus using MM modeling leads to 3-fold increase of RMSd, suggesting structural distortion of $P_i$ from the energetically optimized configuration.

The obtained Ras-GDP-GAP systems were used to perform 106-ps NVE simulations, similarly. Then, we found that 3 of the 49 1-ps NVE simulations for the GDP systems were aborted by 30 fs. This can be due to anomalous configurations generated by the MM-based modeling. The simple MM potential energy function should restrict relocation of the reactive atoms, thus preventing generation of energetically preferable $P_i$ configuration and also GDP (further explanation is given just below). It is noted that the three aborted simulations are exclude for the energy relaxation analyses discussed in **Figure S2**.

As for the 46 snapshot structures derived from the MM modeling, we find that a hydrogen atom in $P_i$ is anomalously close to the phosphate atom, the bond distance between such phosphate-hydrogen pair has 1.33 ± 0.02 Å (**Figure S2A** for representative configuration). An oxygen atom of $P_\beta O_3$ group in GDP also makes anomalously close contact with the oxygen atom of GDP's $P_\alpha O_3$ linked with $P_\beta$, where the distance between such oxygen pair has 1.60 ± 0.07 Å (**Figure S2B** for representative configuration).

These structural distortions reflect on relatively large bonded energy at the switchover timing (see black lines in **Figure S2C, D** and **E**, and compare with **Figure 4A, C** and **D**). The bonded energy of GDP plus $P_i$ for MM modeled Ras-GDP-GAP system is 2.6-fold greater than that for QM/MM modeled Ras-GDP-GAP system. The distorted configurations of GDP and $P_i$ give excessive increase of kinetic energy. The increase from the initial value is 4-fold greater at maximum than the increase discussed in the main text (**Figure S2C**). Accordingly, these structural distortions given by the MM-based modeling definitely prevent us from obtaining physicochemically reasonable observations for mechanical process of GTP-GDP conversion.

It could be supposed that such distortions come from using GTP force field parameters to describe a part of atoms in $P_i$ upon converting GTP to GDP. Thus it may be one technical option to perform additional MM simulations for the modeled Ras-GDP-GAP system with GDP force field parameters in advance of the NVE simulations. Whereas considering such an additional technical step makes the modeling procedure complicated and essentially it is not sure that this post treatment can properly refine artificial structural distortion of configurations of GDP and $P_i$, which is brought about by using the MM-based GDP modeling. Meanwhile, by employing QM/MM simulations, we can straightforwardly obtain snapshot structures which show stable system development and provide physicochemically reasonable observations.

#### SI 4. Angular correlation coefficients analyses over Ras-GAP complex

Angular correlation coefficients were calculated for 488 residues consisting of 166 amino acid (AA) residues in Ras, 320 AA residues in GAP, $Mg^{2+}$ and GTP/GDP. Instead of atomic velocity vectors, we employed mass weighted velocity vectors for each residue, where nonhydrogen atoms are considered.

In the time domain of 0-0.1 ps, the two residue pairs show relatively large difference between Ras-GTP-GAP and Ras-GDP-GAP systems; Ala18-Gln61 (0.060 +/− 0.090 and −0.122 +/− 0.091, respectively), Gly13-$Mg^{2+}$ (−0.002 +/− 0.058 and 0.123 +/− 0.054, respectively). However, these significant differences are not detected in the following time domain, thus seeming to transiently occur due to force field switching and to be irrelevant to the functional expression of Ras.

Focusing on residue pair with magnitude of angular correlation coefficient larger than 0.1, we cannot find any other pair showing significant difference in angular correlation coefficient between Ras-GTP-GAP and Ras-GDP-GAP systems with regard to any time domains. Thus, we conclude that GTP-GDP conversion does not affect global dynamics of Ras-GAP complex within these time scales. As discussed in the main part of this manuscript, GDP and $P_i$ locally interacts with Ras and contributes to change of P-loop in the protein.

### Supporting Tables

**Table S1**. Atomic charges of $H_2PO_4^-$.

| atom name† | atom type† | RESP charge [a.u.] |
|---|---|---|
| O1 | oh | -0.835395 |
| H1 | ho | 0.448002 |
| P1 | p5 | 1.627536 |
| O3 | o | -0.926375 |
| O4 | o | -0.926375 |
| O2 | oh | -0.835395 |
| H2 | ho | 0.448002 |

† Atom name and type follow the style of General Amber Force Field[3].

### Supporting Figures

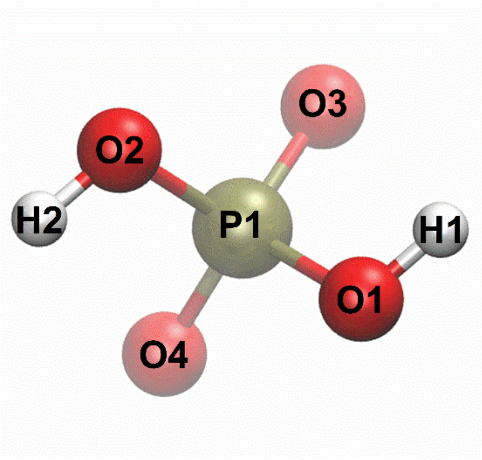


**Figure S1**. Atomic structure of $H_2PO_4^-$, calculated in vacuum with Hartree-Fock/6-31G$^*$ level of theory. Red, white and gold spheres are for hydrogen, oxygen and phosphorus atoms. Characters found inside these spheres denote atom name given as General Amber Force Field style.

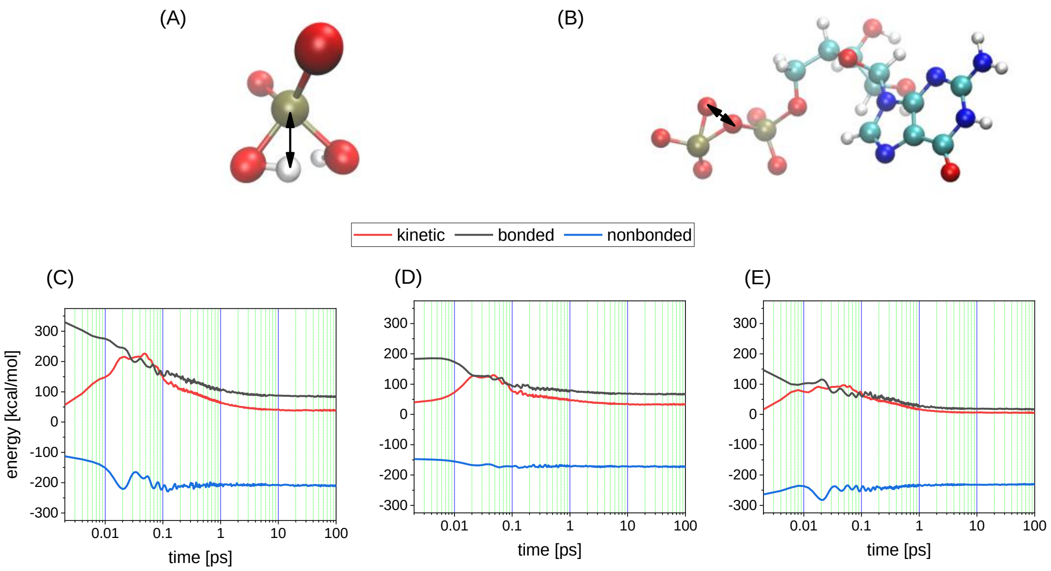


**Figure S2**. Analyses of Ras-GDP-GAP system, modeled by MM simulations. Representative distorted structures of (A) $H_2PO_4^-$ ($P_i$) and (B) GDP. Anomalously closing atom pair is indicated by black double head arrow. The values of distance for the atom pair are 1.33 Å and 1.60 Å in panels A and B, respectively. Time course change of energies for (C) GDP plus $P_i$, (D) GDP and (E) $P_i$. Kinetic, bonded and nonbonded energies are shown with red, black and blue lines, respectively.

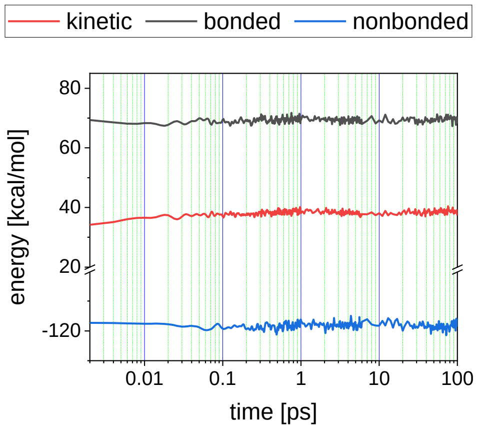


**Figure S3**. Time course analyses of mechanical energy for Ras-GTP-GAP system. Kinetic, bonded and nonbonded energies were shown with red, black and blue lines, respectively. Subtle increase of kinetic energy should result from thermal equilibration of $P\gamma O_3^-$ and the reactive water, which is the counterpart of $H_2PO_4^-$ in the Ras-GDP-GAP system, whose atomic velocities were set to zero at the initial condition.

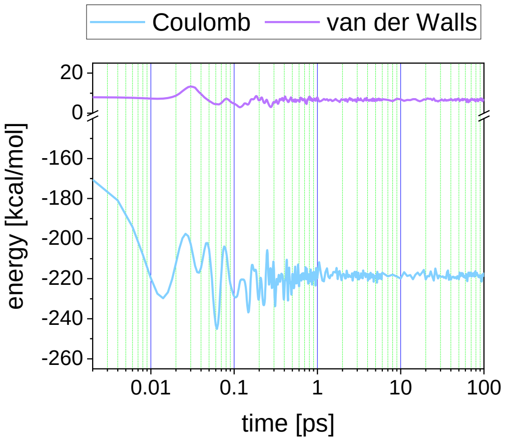


**Figure S4**. Time course analyses of nonbonded energy acting between GDP and $H_2PO_4^-$ in Ras-GDP-GAP system. Coulomb energy and van der Waals energy are individually shown with blue and purple lines, respectively.

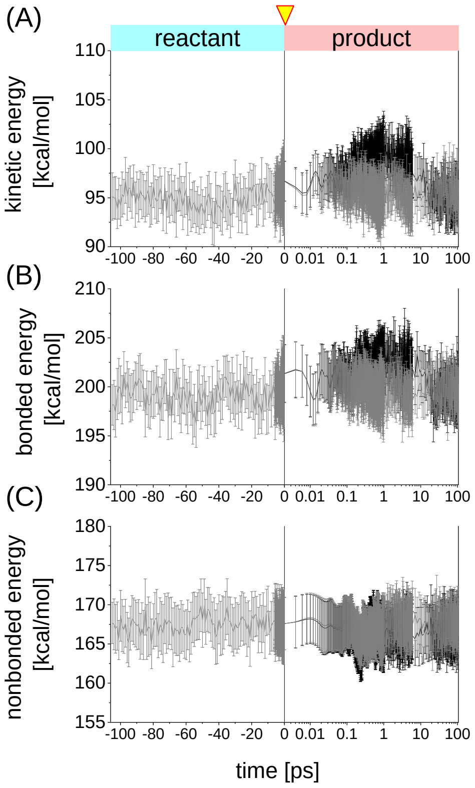


**Figure S5**. Energetic changes of Switch I region in Ras after GTP-GDP conversion. (A)-(C) temporal changes of kinetic, bonded and nonbonded energies, where Ras-GTP-GAP system and Ras-GDP-GAP system are represented by grey and black lines, respectively.

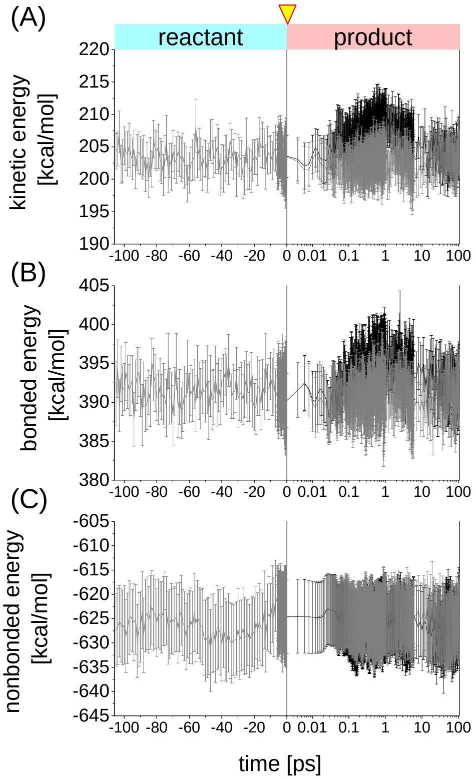


**Figure S6**. Energetic changes of Switch II region in Ras after GTP-GDP conversion. (A)-(C) temporal changes of kinetic, bonded and nonbonded energies, where Ras-GTP-GAP system and Ras-GDP-GAP system are represented by grey and black lines, respectively.

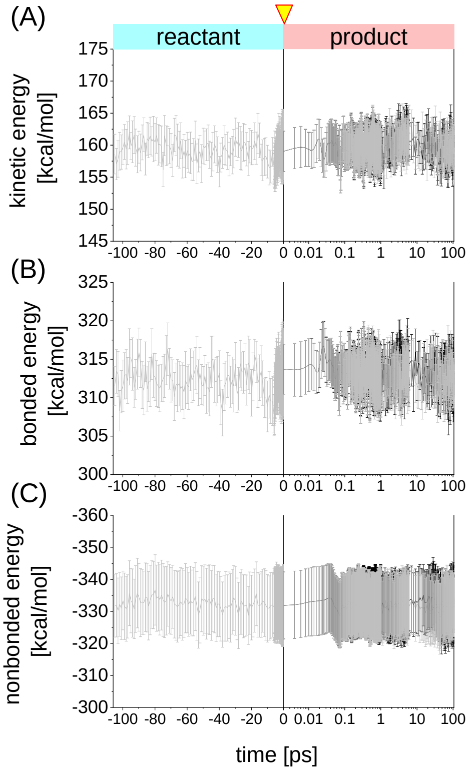


**Figure S7**. Energetic changes of α-helix 3 in Ras after GTP-GDP conversion. (A)-(C) temporal changes of kinetic, bonded and nonbonded energies, where Ras-GTP-GAP system and Ras-GDP-GAP system are represented by grey and black lines, respectively.

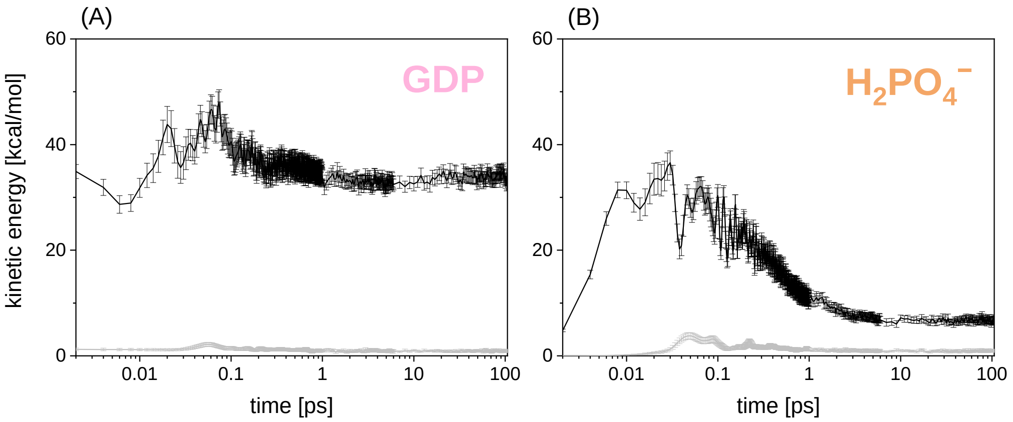


**Figure S8**. Time course analyses of kinetic energy for Ras-GDP-GAP system. (A) GDP and (B) $H_2PO_4^-$. The kinetic energy is decomposed into the contribution from relative motion and that from the center of mass motion (shown by black and gray lines, respectively).

#### References


1. H. J. C. Berendsen, J. P. M. Postma, W. F. Vangunsteren, A. Dinola and J. R. Haak, *J. Chem. Phys.*, 1984, **81**, 3684-3690.
2. K. Ogata, J. W. Shen, S. Sugawa and S. Nakamura, *B. Chem. Soc. Jpn.*, 2012, **85**, 1318-1328.
3. J. M. Wang, R. M. Wolf, J. W. Caldwell, P. A. Kollman and D. A. Case, *J. Comput. Chem.*, 2004, **25**, 1157-1174.
